# Predicting Protein Electrostatics with Protein Language Models

**DOI:** 10.1101/2025.04.17.649309

**Authors:** Mingzhe Shen, Guy W. Dayhoff, Daniel Kortzak, Jana Shen

## Abstract

Ionization states play crucial roles in protein function, yet predicting protein p*K*_a_ values remains a formidable challenge despite decades of research. Here we present KaML-ESM2 and KaML-ESMC, neural network task heads built on ESM protein language models (pLMs) and trained on the PKAD-3r experimental dataset augmented via GAINES, a latent-space sampling strategy for addressing data scarcity. KaML-ESM2/ESMC significantly outperform current structure- and sequence-based approaches across four benchmarks, achieving root-mean-square errors of about 0.5 units across six titratable residue types in native proteins. Performance degradation on 89 buried engineered OBTRUDEs (i**O**nizable su**B**s**T**itutions fo**R** bUrie**D** r**E**sidues) that lack evolutionary support (KaML-ESMC RMSE = 1.89) reveals a key limitation of the current framework, which may be overcome through supervised training. Based on these and other data presented in the work, we hypothesize that protein sequence, through its evolutionary context, encodes not only structure and function but indirectly also electrostatic characteristics. We applied KaML-ESM2 to the human proteome, demonstrating that predicted p*K*_a_ values can potentially identify functional sites and infer catalytic mechanisms. We offer KaML, a sequence-based, end-to-end platform to support applications spanning biological exploration, drug design, protein engineering, and biomolecular simulation. Although additional research is needed, GAINES could offer a general framework for addressing data scarcity in machine learning approaches for protein-related problems.

## Introduction

Ionization states play crucial roles in protein structure and function. In the past, prediction and interpretation of ionization states relied on structure-based approaches, including physics-based^1–3^ and empirical calculations.^4^ Recently, structure-based ML models have been developed and trained on the Poisson-Boltzmann (PB) calculations,^5^ constant pH molecular dynamics (CpHMD) simulations^6,7^ and experimental data.^8–10^ Given that protein functions (e.g., catalysis) are often governed by ionization states, here we hypothesized that the latter can be directly predicted from the protein sequence.

Through masked learning of protein sequences evolved over hundreds of millions of years, evolutionary scale models (ESMs), a class of protein language models (pLMs), demonstrated powerful capacities in predicting structures and functions.^11–13^ This is because representations emerging within the pLM reflect the biological structure and function and improve with scale, e.g., ESM3 is trained with 2.78 billion protein sequences.^12^ We posited that residue-level representations learned by ESMs encode information about residue-specific ionization states and p*K*_a_ shifts that occur when residues transition from solution to protein environment, which are often large for functional sites.

To test the sequence-electrostatics hypothesis, we developed sequence-based ML models, where per-token (i.e., residue-specific) embeddings from the ESM2^11^ or ESMC^13^ model were used as inputs to a feedforward neural network for predicting residue-specific p*K*_a_ values. Training was conducted on a revised experimental p*K*_a_ dataset (PKAD-3r^8^) augmented by synthetic data harvested using a novel approach called GAINES (auGment dAta wIth lateNt spacE Sampling). The new models are named KaML-ESMs, as the training protocol follows our recent development of the structure-based KaML (p***K***_**a**_ **M**achine **L**earning) models, including the tree model KaML-CBT and graph attention neural network KaML-GAT.^8^

Across four benchmark test sets of natural proteins, the KaML-ESM models achieved root-mean-square errors (RMSEs) of about 0.5 units across six amino acid types, significantly outperforming alternative approaches including recent ML models based on structure or sequence, empirical approach, and physics-based method. When testing on the p*K*_a_’s of engineered OBTRUDEs (i**O**nizable su**B**s**T**itutions fo**R** bUrie**D** r**E**sidues) of staphylococcal nuclease (SNase) variants,^14–16^ which lack evolutionary precedent, the RMSE increased substantially, exposing a key limitation of KaML-ESMs, which may be overcome through supervised training in the future. We developed an end-to-end KaML platform and applied KaML-ESM2 to predict the p*K*_a_’s of the entire human proteome, illustrating the potential utility for functional annotation and mechanistic interpretation. We expect the KaML platform to enable a wide range of applications, from understanding biology to supporting drug discovery, protein engineering and MD simulations.

## Results

### ESM representations capture the biochemical environment of protein residues

Before using an ESM model as the foundation for downstream learning and predicting residue-specific protein p*K*_a_’s, we first applied t-distributed stochastic neighbor embedding (t-SNE) to investigate two questions: 1) Do the per-token ESM embeddings (i.e., residue-level representations) distinguish between different titratable amino acids (D, E, C, Y, H, and K)? 2) Do they capture information about residue-specific p*K*_a_ values? We extracted representations of the titratable residues in the experimental p*K*_a_ database PKAD-3r (Methods and Table 3). Two ESM models were employed: ESM2 (trained on ~ 65 million sequences with 650 M parameters)^11^ and the latest ESM Cambrian (ESMC, trained on 2.78 billion sequences with 6 B parameters),^12,13^ which has been demonstrated to predict emergent structures with greater accuracy than ESM2. The t-SNE algorithm^17,18^ was subsequently used to project pairwise residue representation similarities into a two-dimensional space for visualization. The t-SNE maps (Fig. 1a) show that both ESM2 and ESMC distinguish different titratable amino acids, as indicated by the observed clustering patterns. In particular, basic (HK) and acidic (DECY) residues are well separated. However, Asp and Glu residues appear somewhat intermixed in the ESM2 map, which may be attributed to their similar chemical properties and (implicitly) structural environments. Surprisingly, both ESM models fail to distinguish between different types of titratable residues in model peptides or SNase OBTRUDEs (Fig. 1a, circled or boxed regions showing overlapping data). This suggests that the ESM representations capture sequence and structural context, instead of directly encoding the identities of amino acids. It is also note worthy that most SNase OBTRUDEs cluster together despite different sequence positions, suggesting that their shared hydrophobic, buried environment is distinct from those of native ionizable residues (see later discussion).

**Figure 1:**
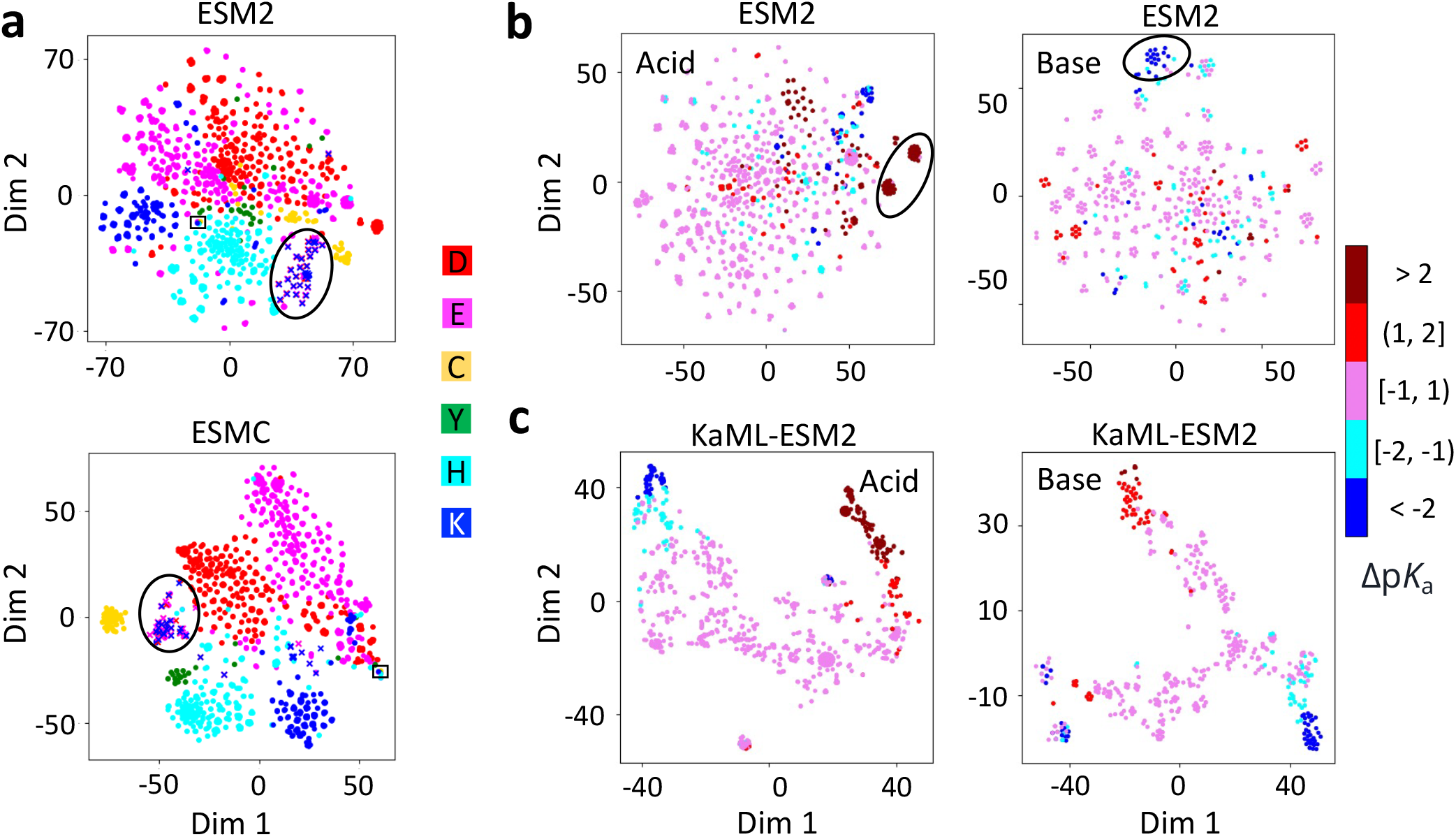
ESM representations can distinguish between titratable amino acids and capture weak signals of residue-specific p*K*_a_ shifts. **a.** t-SNE visualization of the pairwise ESM2 (top) and ESMC (bottom) representation similarities between titratable residues in PKAD-3r dataset. Data points are colored according to amino acid. Circle indicates the SNase OBTRUDEs (in crosses); box indicates model peptide residues. **b, c**. t-SNE visualization of the pairwise representation similarities (ESM2 in b, KaML-ESM2 in c) among acidic (DECY) and basic (HK) residues. Data points are colored according to the shifts of experimental p*K*_a_’s relative to model values. Embeddings were extracted from layer 31 of ESM2_650M (1280-dimensional), layer 80 of ESMC (2560-dimensional), and the final hidden layer of KaML-ESM2 (32-dimensional) trained on PKAD-3r. See Suppl. Fig. S1 for ESMC and KaML-ESMC plots and Fig. S2 for plots without SNase OBTRUDEs.

Next, we asked whether the ESM representations capture p*K*_a_ values by coloring data on the t-SNE map according to their experimental p*K*_a_ shifts relative to the (solution) model values. While most data points with positive and negative p*K*_a_ shifts are mixed, a few clusters with p*K*_a_ shifts greater than 1 are visible for both acids and bases (Fig. 1b and Fig. S1a for ESMC analysis). Comparison of the maps with and without the SNase OBTRUDEs reveals that the two largest acid clusters of upshifted p*K*_a_’s and the largest base cluster of downshifted p*K*_a_’s belong to the SNase OBTRUDEs (Fig. 1b and Fig. S2), which reinforces the notion that OBTRUDEs and native titratable residues have dissimilar environments. Collectively, the t-SNE analysis demonstrates that while the ESM representations capture some weak p*K*_a_ signals, they cannot be directly used for p*K*_a_ predictions. We will come back to this point.

### Developing the KaML-ESM task heads for protein p*K*_a_ prediction

Having assessed the suitability of ESM2 or ESMC as foundation model, we proceeded to building a four-layer multilayer perceptron (MLP) task head for protein p*K*_a_ predictions. The MLP task head utilizes the ESM2 or ESMC representations of the input sequence, and model training was performed on the experimental p*K*_a_ shifts from the PKAD-3r database. Separate MLPs were trained for acidic and basic residues, following our previous work.^8^ We evaluated whether fine-tuning of ESM2_650M improved performance relative to training task heads on frozen embeddings. Using the parameter-efficient finetuning method qLoRA^19^ (Suppl. Methods), fine-tuning decreased RMSE for some amino acids but increased RMSE for others, resulting in a negligible overall RMSE decrease of 0.04 (Suppl. Table S6). Due to the mixed performance and substantial increase in computational cost, we chose to not to use finetuning for the remainder of this work.

To avoid potential overfitting, acid and base models were pretrained on a synthetic dataset comprising 29,457 p*K*_a_ shifts of 9,945 proteins from the PDB predicted by the structurebased KaML-CBT model^8^ (Suppl. Methods and Fig. S3). The pretrained models were then trained on the experimental PKAD-3r dataset (Fig. S4). Both pretraining and separating acid and base models decreased the p*K*_a_ errors (Suppl. Table S2). Since protein p*K*_a_ shifts are primarily determined by local structure environment^8,20^ and previous work found that local contacts are learned across different transformer layers in the pLM,^21^ we analyzed different ESM layers by training models using each layer’s embeddings. The second-to-last layer of ESM2 and the last layer of ESMC gave the lowest overall test RMSEs; therefore, these layers were selected for further development. We also examined the potential impact of increasing ESM’s parameter scale by comparing models trained with ESM2 used so far (ESM2_650M) with a larger ESM2 (ESM2_15B). Surprisingly, KaML-ESM2_15B gave a larger overall RMSE and was therefore excluded from further discussion. Details are given in Methods, Suppl. Table S3-S5 and Fig. S5.

To disentangle the contributions of the ESM foundation model and task-specific training, we performed ablation studies using a baseline model KaML-ESM2 (without pretraining on the KaML-CBT predicted p*K*_a_’s), which gave an RMSE of 0.75 across 50 stratified random holdout sets. The zero-shot predictions and a task head trained for a single epoch gave average RMSEs of 1.59 and 0.90, respectively, confirming that the ESM2 representations alone are insufficient and task-specific training is required for p*K*_a_ predictions, consistent with the t-SNE analysis (Fig. 1b and c). A KaML task head trained using the first-layer ESM2 representations yielded an RMSE of 1.22, compared to 0.75 using the second-to-last layer presentations (baseline), demonstrating that ESM2’s learning of protein evolution generates protonation state-relevant information. We also trained a KaML task head on ProstT5,^22^ another recent pLM, which gave an RMSE of 0.89, supporting the choice of ESM2 as the foundation. Finally, KaML-ESM2 trained on randomly permuted p*K*_a_ labels gave an RMSE of 1.37. Detailed metrics of ablation experiments are given in Suppl. Table S8.

## Developing GAINES to overcome data scarcity

We examine the performance of KaML-ESM2/ESMC (with pretraining) across 50 stratified random holdout sets (Table 1). The average overall RMSEs of KaML-ESM2 and KaML-ESMC are 0.70 and 0.66, respectively, which are about 0.8 unit lower than the corresponding zero-shot predictions. The RMSEs for Asp, Glu, His, and Lys are 0.61–0.69 with KaML-ESM2 and 0.53–0.66 with KaML-ESMC, while the RMSEs for Cys and Tyr are much larger, 1.1/1.2 and 1.0/1.2, respectively. The poor performance for Cys and Tyr reflects extremely limited training data (60 and 39 p*K*_a_ values, respectively), exemplifying the data scarcity challenge common in scientific ML.

**Table 1:** Performance of the GAINES-trained KaML-ESM models in comparison to conventionally trained models and zero-shot predictions.

|  |  | KaML-ESM2 |  |  | KaML-ESMC |  |  |
| --- | --- | --- | --- | --- | --- | --- | --- |
|  |  | RMSE | PCC | CER | RMSE | PCC | CER |
| Asp | 0-shot | $1.80 \pm 0.03$ | $0.02 \pm 0.03$ | 148/2091 | $1.65 \pm 0.05$ | $0.08 \pm 0.05$ | 135/2089 |
| | Conv | $0.65 \pm 0.04$ | $0.91 \pm 0.01$ | 7/2018 | $0.66 \pm 0.03$ | $0.90 \pm 0.01$ | 17/2028 |
| | **GAINES** | **$0.54 \pm 0.02$** | $0.95 \pm 0.01$ | 8/1992 | $0.52 \pm 0.04$ | $0.94 \pm 0.01$ | 18/2024 |
| Glu | 0-shot | $1.46 \pm 0.03$ | $-0.02 \pm 0.02$ | 94/2431 | $1.30 \pm 0.04$ | $0.08 \pm 0.04$ | 88/2446 |
| | Conv | $0.61 \pm 0.04$ | $0.85 \pm 0.02$ | 10/2418 | $0.62 \pm 0.02$ | $0.83 \pm 0.02$ | 38/2433 |
| | **GAINES** | **$0.42 \pm 0.02$** | $0.94 \pm 0.01$ | 8/2405 | $0.56 \pm 0.04$ | $0.87 \pm 0.02$ | 31/2426 |
| Cys | 0-shot | $2.96 \pm 0.15$ | $-0.04 \pm 0.09$ | 87/174 | $2.75 \pm 0.15$ | $-0.10 \pm 0.10$ | 79/172 |
| | Conv | $1.11 \pm 0.10$ | $0.74 \pm 0.11$ | 34/173 | $1.19 \pm 0.09$ | $0.68 \pm 0.11$ | 24/167 |
| | **GAINES** | **$0.50 \pm 0.05$** | $0.95 \pm 0.02$ | 4/178 | $0.57 \pm 0.08$ | $0.95 \pm 0.02$ | 5/178 |
| Tyr | 0-shot | $1.15 \pm 0.11$ | $-0.13 \pm 0.12$ | 0/85 | $1.33 \pm 0.12$ | $0.07 \pm 0.12$ | 4/96 |
| | Conv | $1.18 \pm 0.11$ | $-0.13 \pm 0.12$ | 4/96 | $0.98 \pm 0.08$ | $0.32 \pm 0.21$ | 4/99 |
| | **GAINES** | **$0.32 \pm 0.03$** | $0.34 \pm 0.11$ | 0/88 | $0.92 \pm 0.16$ | $0.34 \pm 0.21$ | 4/96 |
| His | 0-shot | $1.25 \pm 0.03$ | $0.03 \pm 0.03$ | 170/515 | $1.02 \pm 0.03$ | $0.00 \pm 0.04$ | 76/360 |
| | Conv | $0.68 \pm 0.03$ | $0.69 \pm 0.04$ | 19/565 | $0.57 \pm 0.02$ | $0.80 \pm 0.03$ | 5/665 |
| | **GAINES** | **$0.39 \pm 0.02$** | $0.92 \pm 0.01$ | 16/751 | $0.37 \pm 0.02$ | $0.93 \pm 0.01$ | 6/769 |
| Lys | 0-shot | $1.40 \pm 0.05$ | $-0.07 \pm 0.04$ | 23/776 | $1.13 \pm 0.04$ | $0.05 \pm 0.05$ | 21/775 |
| | Conv | $0.69 \pm 0.07$ | $0.78 \pm 0.04$ | 14/770 | $0.53 \pm 0.02$ | $0.88 \pm 0.02$ | 8/756 |
| | **GAINES** | **$0.52 \pm 0.03$** | $0.89 \pm 0.01$ | 13/769 | $0.45 \pm 0.03$ | $0.90 \pm 0.03$ | 8/765 |
| **All** | 0-shot | $1.59 \pm 0.02$ | $0.78 \pm 0.01$ | 522/6072 | $1.42 \pm 0.03$ | $0.82 \pm 0.01$ | 403/5938 |
| | Conv | $0.70 \pm 0.02$ | $0.96 \pm 0.01$ | 88/6040 | $0.66 \pm 0.02$ | $0.96 \pm 0.01$ | 96/6145 |
|  | **GAINES** | **$0.48 \pm 0.02$** | **$0.98 \pm 0.01$** | **49/6183** | **$0.53 \pm 0.02$** | **$0.97 \pm 0.01$** | **72/6258** |

The zero-shot predictions from the ESM models are compared to those from the KaML-ESM models trained using either the conventional (Conv) or the GAINES augmentation protocol. The average and standard error of RMSEs, Pearson correlation coefficients (PCC), and critical error rates (CERs), computed over 50 stratified random holdout sets of PKAD-3r are listed. CER refers to the fraction of misclassified protonation states at pH 7.<sup>8</sup>

To tackle data scarcity challenge, we developed a data augmentation strategy called GAINES (auGment dAta wIth lateNt spacE Sampling). Inspired by the attention mechanism in transformers,^23^ GAINES uses training data as queries to retrieve data from an unrelated database based on embedding similarity. The retrieved data are then assigned the labels of their corresponding training queries. We implemented a GAINES protocol for augmenting experiment p*K*_a_ dataset (Fig. 2a). For each query residue (e.g., a cysteine with an experimental p*K*_a_), its ESM2 or ESMC representation serves as the query embedding, which is compared against the embeddings of all residues (keys) in a protein database (proteins overlapping the training set excluded). If the cosine similarity between query and key embeddings exceeds a predefined threshold (e.g., 0.8 used in this work), the value residue is assigned the query residue’s p*K*_a_ and added to the synthetic data pool. For convenience in testing structure-based KaML-CBT models, we used RCSB PDB as the protein database in this work. The GAINES protocol yielded a large pool of synthetic p*K*_a_ data; however, to preserve the p*K*_a_ distributions similar to the experimental PKAD-3r dataset, we randomly down-sampled to a maximum of 10 value residues (and proteins) per query residue. The resulting augmented dataset comprises p*K*_a_ data of 4,220 unique residues (1,055 Asp, 1,209 Glu, 756 His, 675 Cys, 417 Lys, and 108 Tyr) from 2,164 proteins, approximately ten times the experimental dataset (Suppl. Fig. S7 and Table S1).

**Figure 2:**
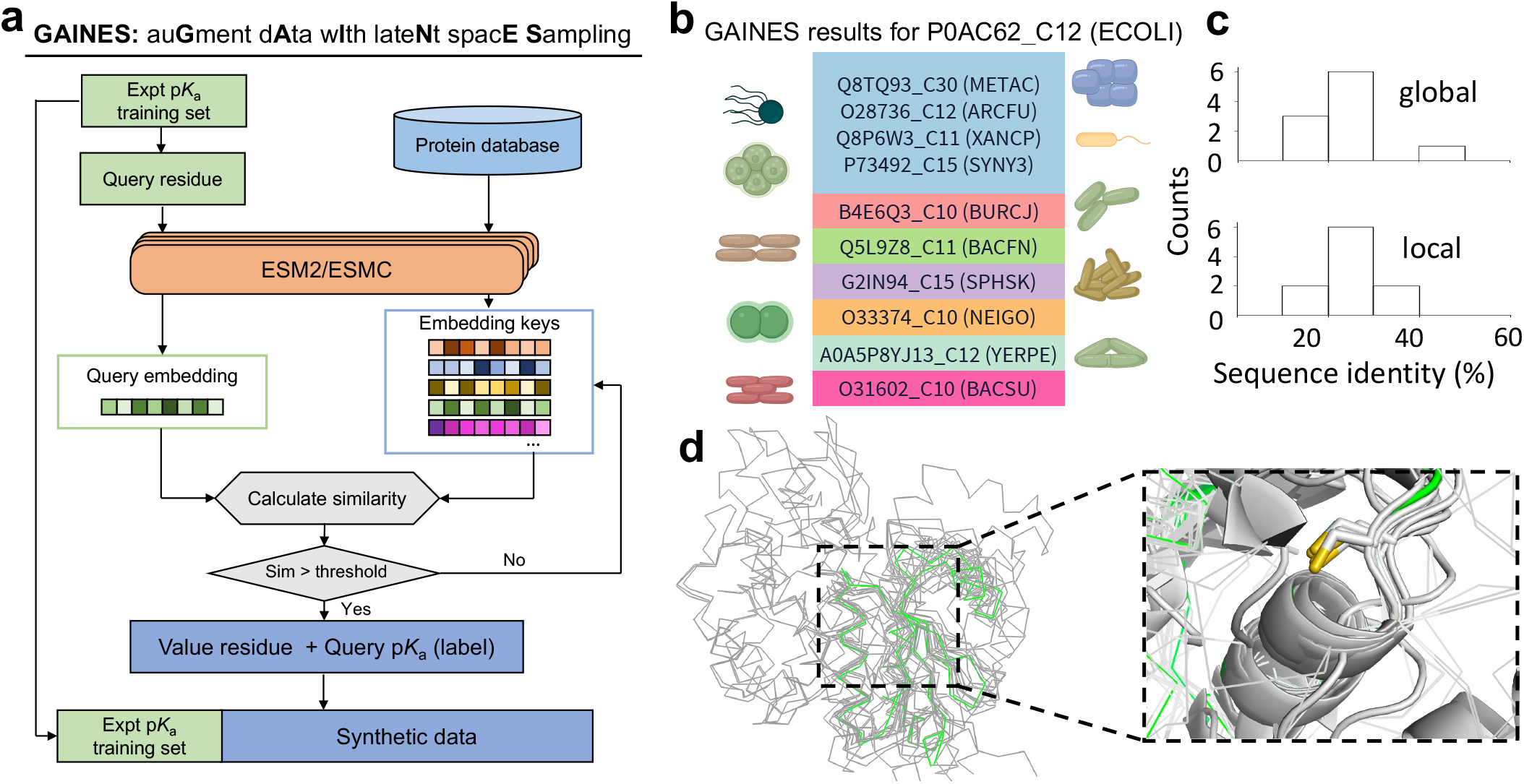
GAINES protocol and its application to identify synthetic data related to a downshifted cysteine p*K*_a_ in glutaredoxin 3. **a.** The GAINES flowchart. The query, key, and value concept is used to identify synthetic data from an external database. **b**. GAINES identified 10 value proteins across 10 species (in parenthesis) that contain a cysteine similar to query C12 in E. Coli glutaredoxin 3 (UniProt ID: P0AC62). **c**. A histogram of the global and local sequence identities between query (P0AC62) and value proteins. **d**. Overlay of the ESM3-predicted structures of P0AC62 (green) and value proteins (gray). A zoomed-in view shows a similar local environment (within 5 Å) for the query and value cysteines (yellow and green sticks).

A potential concern is that residue embedding similarity may simply reflect sequence similarity rather than meaningful structural or physicochemical relationships, which could hinder model generalization. This is however not the case, as the majority of the query and value protein pairs share a sequence identity below 40% (Suppl. Fig. S8). Taking Cys12 in E. Coli glutaredoxin 3 (experimental p*K*_a_ of 4.1^24^) as an example, the global and local sequence (within 10 amino acids from the cysteine of interest) identities between P0AC62 and most (9 out of 10) value proteins (from different species) fall below 40% (Fig. 2b and 2c). The AlphaFold2-generated structures demonstrate that while global structures may differ, the local environment around the cysteine of interest is highly similar (Fig. 2d). Protein family analysis with Pfam^25^ reveals that all these proteins are related as glutathione S-transferase-dependent enzymes, suggesting that embedding similarity captures functional conservation across evolutionarily distant species.

Using the GAINES-augmented training set, we retrained KaML-ESM2/ESMC and tested on the aforementioned 50 holdout sets (no synthetic data, Supplemental Methods). Strikingly, the RMSEs for all residue types are decreased for both KaML-ESM2 and KaML-ESMC (Table 1 and Suppl. Table S7). As expected, the most significant performance gain is for Cys and Tyr p*K*_a_’s, with the RMSE of KaML-ESM2 decreased to 0.50 and 0.32, respectively. Note, the latter needs to be taken with a grain of salt due to the extremely small test set. Similarly, the RMSEs of KaML-ESMC for Cys and Tyr p*K*_a_’s are decreased to 0.57 and 0.92, respectively. Importantly, the critical error rate, defined as the fraction of misclassified protonation states at pH 7, is reduced by 8 or 5 times to 4 or 5 out of 178 for Cys for KaML-ESM2 or KaML-ESMC, respectively (Table 1). We also tested the models with and without GAINES training on EXP67S,^6^ a challenging holdout set comprising 160 p*K*_a_’s of Asp, Glu, His, and Lys residues in 36 proteins. The overall RMSEs of KaML-ESM2 and KaML-ESMC are reduced by 0.35 and 0.23, respectively (Suppl. Data File).

To demonstrate that the model performance gains stem from meaningful augmentation rather than increased data volume, we duplicated the training data and increased it 10-fold. The resulting model performed identically to the conventionally trained model without GAINES (Fig. 3). We also tested whether latent space similarity is a critical requirement by varying the embedding similarity used to select synthetic data. The similarity ranges of 0.2–0.4 and 0.4–0.6 increased the model RMSE by 0.20 and 0.07 units, respectively, compared to the conventionally trained model, while the similarity range of 0.6–0.8 resulted in a similar RMSE. (Fig. 3). When the similarity requirement was raised to *>*0.8, the model RMSE was decreased by 0.22 units compared to conventionally trained model (Fig. 3). This trend suggests that assigning query residue labels to value residues from dissimilar biochemical environments introduce noise and degrade model performance, whereas restricting label assignment to value residues from similar environments yields meaningful augmentation.

**Figure 3:**
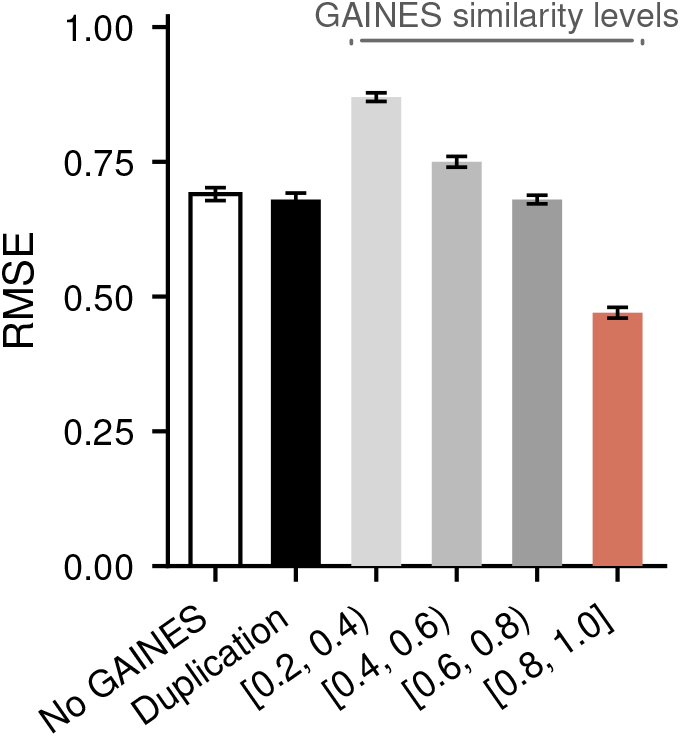
Ablation of the GAINES strategy. RMSEs of trial KaML-ESM2 models on 50 stratified random holdout sets. Models were trained without GAINES, with a 10-fold duplicated training set, or with GAINES-augmented data generated at varying levels of embedding similarities.

To test whether the GAINES training can also boost the performance of structure-based p*K*_a_ models, we retrained KaML-CBT using a GAINES augmented training set, in which protein structures corresponding to value residues and sequences were added.^8^ GAINES significantly lowers the RMSEs for predicting p*K*_a_’s of Cys, His, Tyr, and Lys, whereas the RMSE decreases for Asp and Glu are not statistically significant (Suppl. Table S7). Since Asp and Glu have the largest training sets, this confirms that GAINES is most effective when data are scarce. We note that GAINES requires a strong pLM that accurately encodes biochemical environments; an insufficiently precise model will introduce excessive noise into training and degrade task head performance. Additionally, the optimal similarity threshold for p*K*_a_ augmentation may not generalize to other tasks and should be tuned accordingly.

### Further benchmarking the KaML-ESM models

#### CD-HIT-partitioned evaluation

To examine whether model performance evaluated by holdouts is inflated by local sequence similarity among a small number of residues in homologous proteins (Suppl. Fig. S9), we conducted two experiments. First, we reevaluated the 50 random holdouts by removing holdout residues with local sequence identity above 40% with any one in the corresponding training set. The local sequence is defined as a 20-residue window around the titratable residue of interest. The resulting overall RMSE is 0.48±0.03 (detailed metrics in Suppl. Data file). Second, we generated 20 splits of the PKAD-3r dataset, using the CD-HIT program^26^ with a local sequence identity threshold of 70% (Suppl. Fig. S10). Note, due to mutual sequence identities SNase OBTRUDEs were excluded and will be separately evaluated (see later discussion). The resulting RMSE is 0.43±0.02 and 0.40 ± 0.02 after additionally imposing a 40% local sequence identity filter (detailed metrics in Suppl. Data file).

#### The EXP67S benchmark

We tested whether the 50 holdout performance extends to the more balanced EXP67S benchmark,^6^ which includes large p*K*_a_ shifts (among a total of 160 p*K*_a_ values in 36 unique proteins after cleaning) and has been used for comparison by previous studies.^6,7,10,27^ KaML-ESM2/ESMC retrained after excluding EXP67S achieve overall RMSEs of 0.52 and 0.56, respectively, comparable to their random holdout performance, confirming their generalizability (Table 2).

**Table 2:**
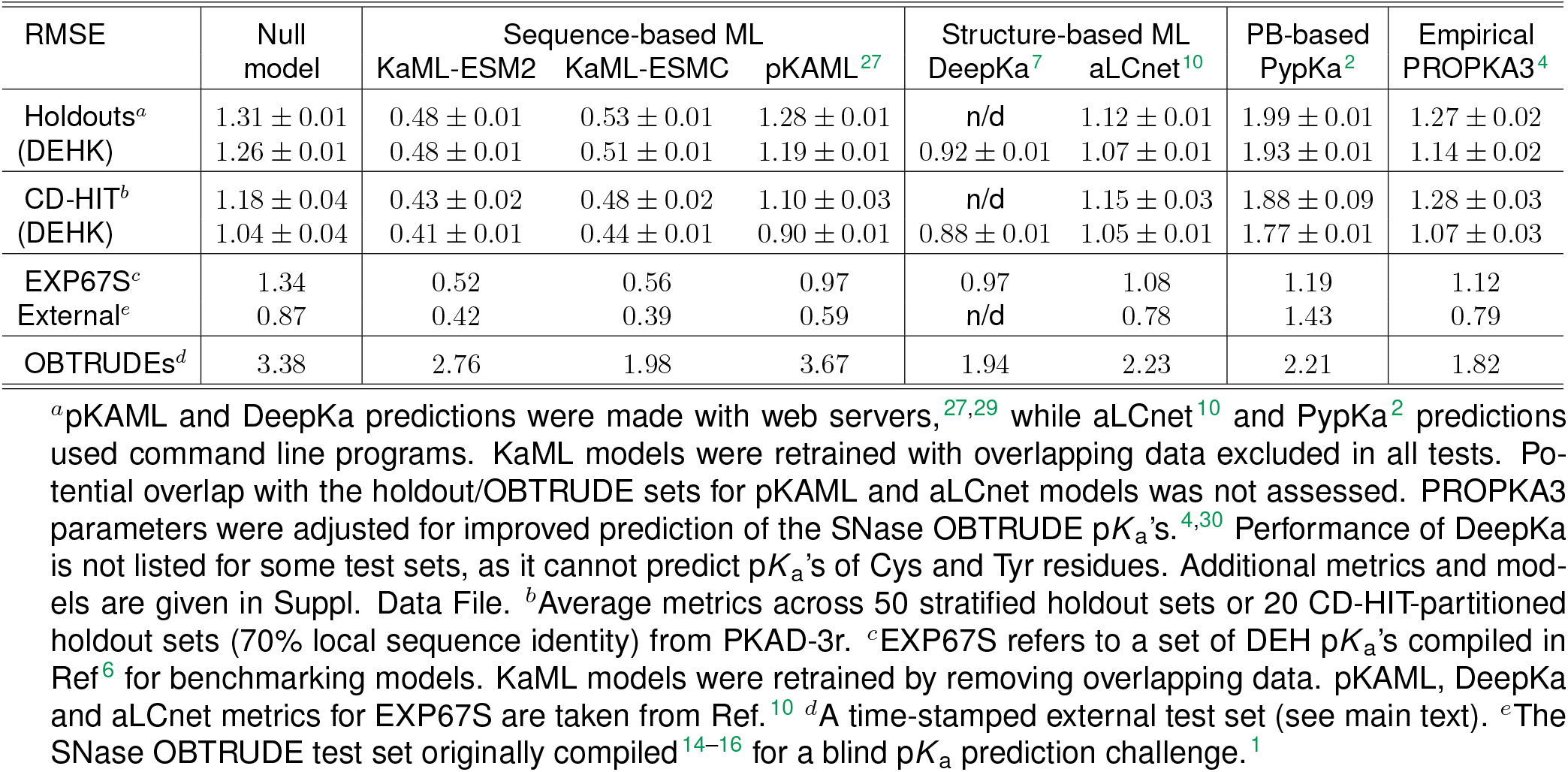
Benchmarking KaML-ESM models against alternative approaches for p*K*_a_ prediction^*a*^.

**Table 3:**
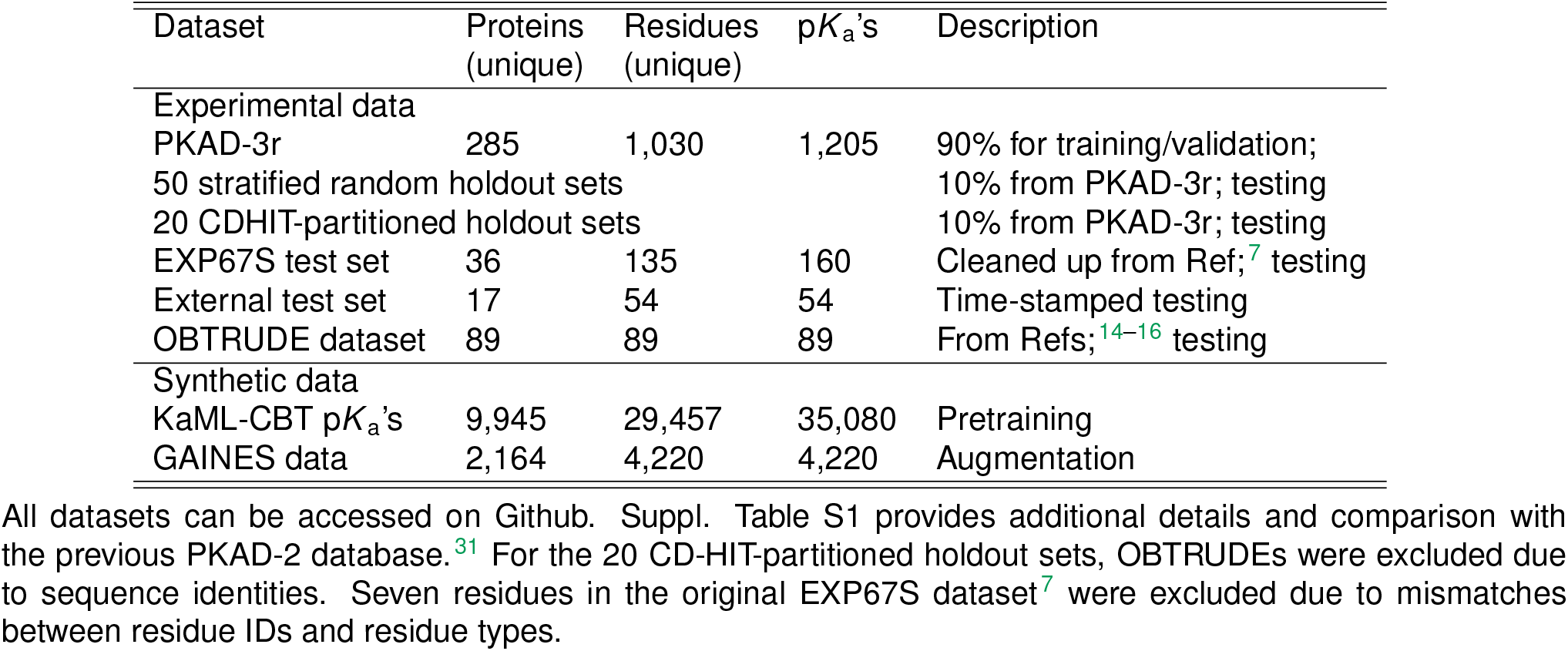
Summary of the p*K*_a_ datasets used in this work^*a*^.

#### Time-stamped external evaluation

We further evaluated KaML-ESMs on experimental p*K*_a_ data published after April 2023 (the PKAD-3 timestamp^8^) with no overlap with the training set. On this external test set of 54 p*K*_a_ values (39 His, 1 Cys, 1 Tyr, and 13 Lys) from 16 proteins, KaML-ESM2 and KaML-ESMC achieve overall RMSEs of 0.42 and 0.39, respectively (Supplemental Data File).

#### The engineered SNase OBTRUDEs

To understand the model weakness, we analyzed the outliers in the 50 holdout tests (Suppl. Fig. S11 and Fig. S12). Unsurprisingly, the majority of them are associated with the engineered SNase OBTRUDEs. We trained KaML-ESM2_no_OB and KaML-ESMC_no_OB models with SNase OBTRUDEs excluded from training, which gave RMSEs of 2.76 and 1.98, respectively, substantially larger than native residues (Table 2) and only somewhat smaller than the zero-shot predictions (3.50 with ESM2 and 3.32 with ESMC). Notably, the p*K*_a_ prediction errors for Lys (RMSE=1.52/1.42) are much lower than those for Asp (RMSE=3.54/2.06) and Glu (RMSE=3.03/2.36, Suppl. Data File). The weak performance for the engineered OBTRUDEs is expected, because their (artificial) environments corresponding to hydrophobic parent residues differ markedly from those of natural titratable residues (Fig. 1b and Suppl. Fig. S2). Consistent with this notion, the GAINES protocol did not detect any comparable residues in the PDB, suggesting that the engineered OBTRUDEs represent environments not observed in nature for titratable sites.

The distinct environments of OBTRUDEs suggest that supervised training is necessary for improving predictions of their p*K*_a_’s. To test this, we focused on two mutation sites, V66 and V74, which exhibit large p*K*_a_ prediction errors. The KaML-ESM2_no_OB predictions for V66D/E exhibit errors greater than 2, while those for V74D/E show errors exceeding 4 (Fig. 4). We retrained the model including only OBTRUDEs located more than 10 amino acids from the mutation site. Errors for V66D/E and V74D/E are substantially reduced, while the error for V66K increases and that for V74K decreases. The overall RMSE of the six OBTRUDEs is 1.61, compared to 3.15 with all OBTRUDEs excluded in training (Fig. 4). This ablation study supports our hypothesis that the MLP task head can learn from the representation–p*K*_a_ relationships of some OBTRUDEs and generalize to others.

**Figure 4:**
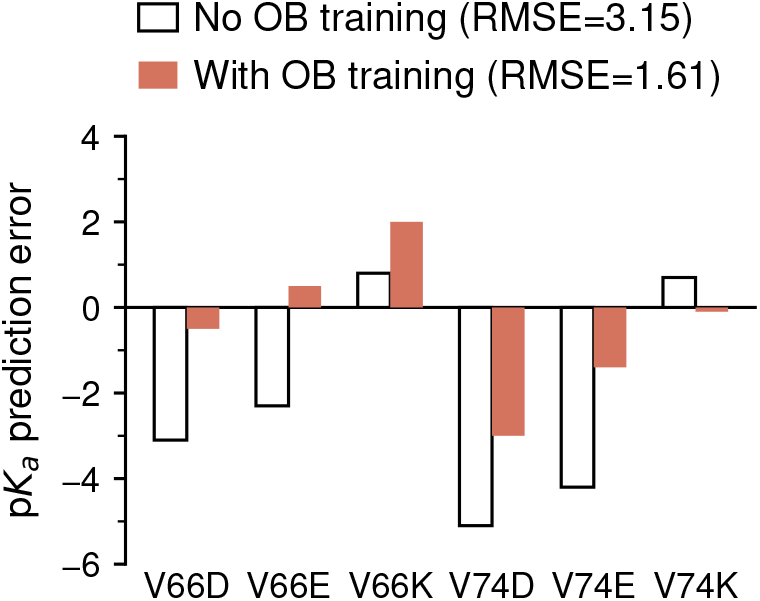
Training improves p*K*_a_ prediction for SNase OBTRUDEs. The KaML-ESM2_no_OB model gave large errors in predicting the p*K*_a_’s of V66D, V66E, V74D, and V74E (empty bars). Models retrained by including OBTRUDEs more than 10 residues from test residue (V66 or V74) gave significantly reduced errors. The RMSE also includes the p*K*_a_ predictions for V66K and V74K.

### Comparison against alternative approaches

We compare the performance of KaML-ESM2/ESMC models against ten alternative approaches. They include the baseline null model (assuming model p*K*_a_ values), the Poisson-Boltzmann (PB) method PypKa,^2^ the empirical method PROPKA3,^4^ and six ML models: pkaani,^9^ aLCnet,^10^ KaML-CBT/CBT2,^8^ pKAML,^27^ pKAI/pKAI+,^5^ and DeepKa.^7^ The first four ML models were trained on experimental p*K*_a_’s^8–10,27^ while the latter two models^5,7^ were trained on p*K*_a_’s calculated by the PB solver^2^ or GBNeck2-CpHMD simulations.^28^ Detailed metrics are provided in the Supplemental Data File. We note that the model comparison here focuses on performance and does not account for differences in training data, as we are unable to retrain the alternative ML models.

Across the stratified and CD-HIT-partitioned holdout sets, EXP67S, and the external test sets, the KaML-ESM2/ESMC models achieve RMSE values around 0.5 pH units, significantly outperforming the null model, other sequence- and structure-based ML approaches, the PB solver, and the empirical method (Table 2 and Suppl. Data File). Among the alternative approaches, the structure-based convolutional neural network (CNN) DeepKa,^7^ which was trained on 26,552 p*K*_a_ values of DEHK residues (549 PDB structures) computed with GBNeck2-CpHMD titration simulations,^28^ is the second-best performer, with RMSE values just under 1 unit. The recently developed graph neural network (GNN) aLCnet^10^ yields RMSEs of roughly 1.1. Surprisingly, the RMSEs of the widely used empirical method PROPKA3^4^ are comparable to those of the null model, and those of the PB solver (PypKa)^2^ are much larger. Across both the stratified and the CD-HIT-partitioned holdout test sets, KaML-ESMs exhibited low critical error rates (i.e., misclassified protonation states), below 1.2% and 0.6%, respectively, which are four times lower than DeepKa.

Among the ML approaches, pKAML^27^ is the only other model that operates on sequences, and it is likewise derived from ESM2. However, its RMSEs across the stratified and CD-HIT-partitioned holdout sets are at or above 1.1 units, and the RMSE on the EXP67S test is close to 1 unit (Table 2). This may be attributed to several key differences in model architecture and training.^27^ pKAML uses a single fully connected layer that takes the concatenated embeddings from ESM2 and those generated by a protein isoelectric point predictor. Moreover, pKAML employs a single model to predict both acidic and basic p*K*_a_ values and does not incorporate any pretraining or synthetic data augmentation. Lastly, pKAML was trained on the PKAD-2 database^31^ by splitting the training and holdout sets in a 5:2 ratio.

Relative to the native residues, every alternative method performs significantly worse for the SNase OBTRUDEs, which exhibit a null-model RMSE of 3.38 as a consequence of large p*K*_a_ shifts. Although the parameters of PROPKA3^4^ were optimized to better match the experimental p*K*_a_ values of SNase OBTRUDEs,^30^ the RMSE remains relatively high at 1.82, which is only about 0.1 unit lower than the RMSEs of DeepKa (1.94) and KaML-ESMC (1.98). Note, the critical error rate of DeepKa is 18% lower than KaML-ESMC. Nonetheless, all of the methods appear to have a higher RMSE than the temperature replica-exchange implicit-solvent GBSW-CpHMD^32,33^ titration simulations, which reported an RMSE of 1.69 for 87 (out of a total of 89) SNase OBTRUDEs^34^ in the blind p*K*_a_ prediction challenge in 2011.^1^

### A KaML platform for p*K*_a_ predictions and visualization

To further extend the applicability of KaML-ESMs, we created a KaML platform that additionally leverages ESM3^12^ to generate protein structures, enabling visualization and, if desired, structure-based p*K*_a_ predictions using KaML-CBT2 (Fig. 5 left, Suppl. Methods). Both a command-line program and a browser-based GUI (https://kaml.computchem.org) are provided, which support input via protein sequence, UniProt ID, PDB ID, or user-provided PDB file (Suppl. Methods and Suppl. Fig. S14). As an example, KaML-ESM2 was applied to predict p*K*_a_’s of UCHL1 (Fig. 5 right), which belongs to the ubiquitin hydrolase family featuring a conserved Cys-His-Asp catalytic triad. KaML-ESM2 predicted p*K*_a_ of 4.67±0.03 for Cys90, 6.95 ± 0.02 for His161, and 2.37± 0.02 for Asp176, allowing functional and mechanistic interpretations (see later discussion).

**Figure 5:**
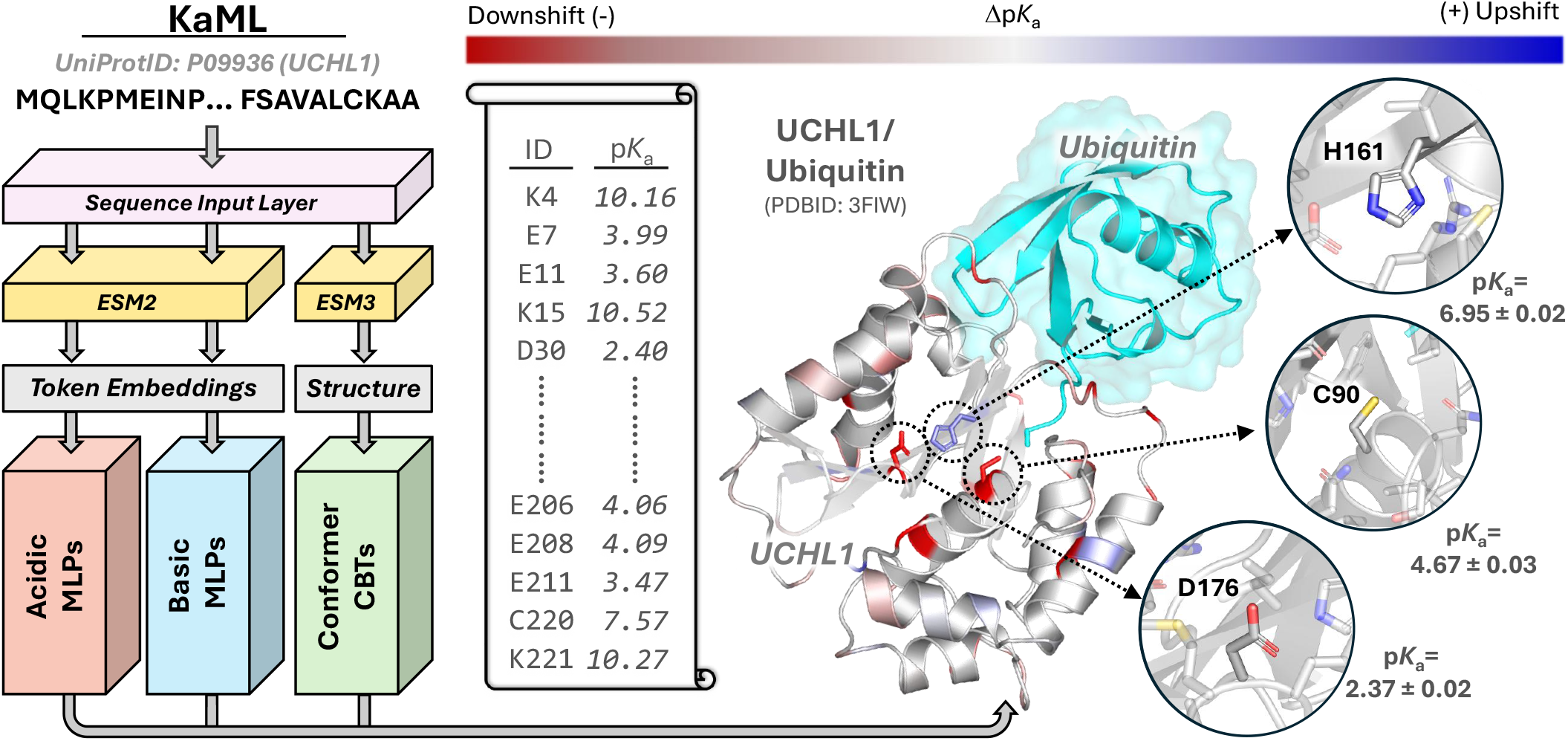
Architecture of the KaML platform and illustration of p*K*_a_ predictions. KaML accepts a user-provided protein sequence through the sequence input layer. ESM2 (or ESMC) is used to generate token embeddings, while ESM3 predicts a three-dimensional structure if not provided. Embeddings are fed to the acid and base task heads (ensemble of 500 MLPs), respectively. Optionally, the ESM3-derived or user-provided structure is processed for conformer-dependent predictions by KaMLs-CBT2. The predicted acid and base p*K*_a_’s, standard errors, and p*K*_a_ shifts are given in the output. A vertical scroll illustrates the output for a human deubiquitinase UCHL1. A cartoon representation of UCHL1 is given, with substrate ubiquitin (not included in the prediction) shown in cyan. Residues with up- and down-shifted p*K*_a_’s are colored in blue and red, respectively. The catalytic triad and their KaML-ESM2 predicted p*K*_a_’s are highlighted and discussed in main text. The inferred catalytic mechanism is displayed in Fig. 6c.

### Proof-of-concept biological applications of KaML-ESM predictions

#### p*K*_a_ prediction for the entire human proteome

Knowledge of protein p*K*_a_ values is valuable for functional and mechanistic studies. We applied KaML-ESM2 to the human proteome, which resulted in 1,877,704 p*K*_a_ values of Asp, Glu, His, Cys, Tyr, and Lys residues in 18,192 proteins with unique genes (Fig. 6a). Note, 1,882 proteins were ex-cluded due to sequence lengths exceeding the limit for ESM2 (1022). This application is not feasible with structure-based models, as most targets lack experimental structures and AlphaFold predictions^37^ contain large low-confidence regions. Interestingly, the median predicted p*K*_a_’s for all residue types are within 0.3–0.4 units from the corresponding solution model p*K*_a_’s (Fig. 6a), even though they were not used as model constraints. This consistency also suggests the absence of systematic errors, at least superficially.

**Figure 6:**
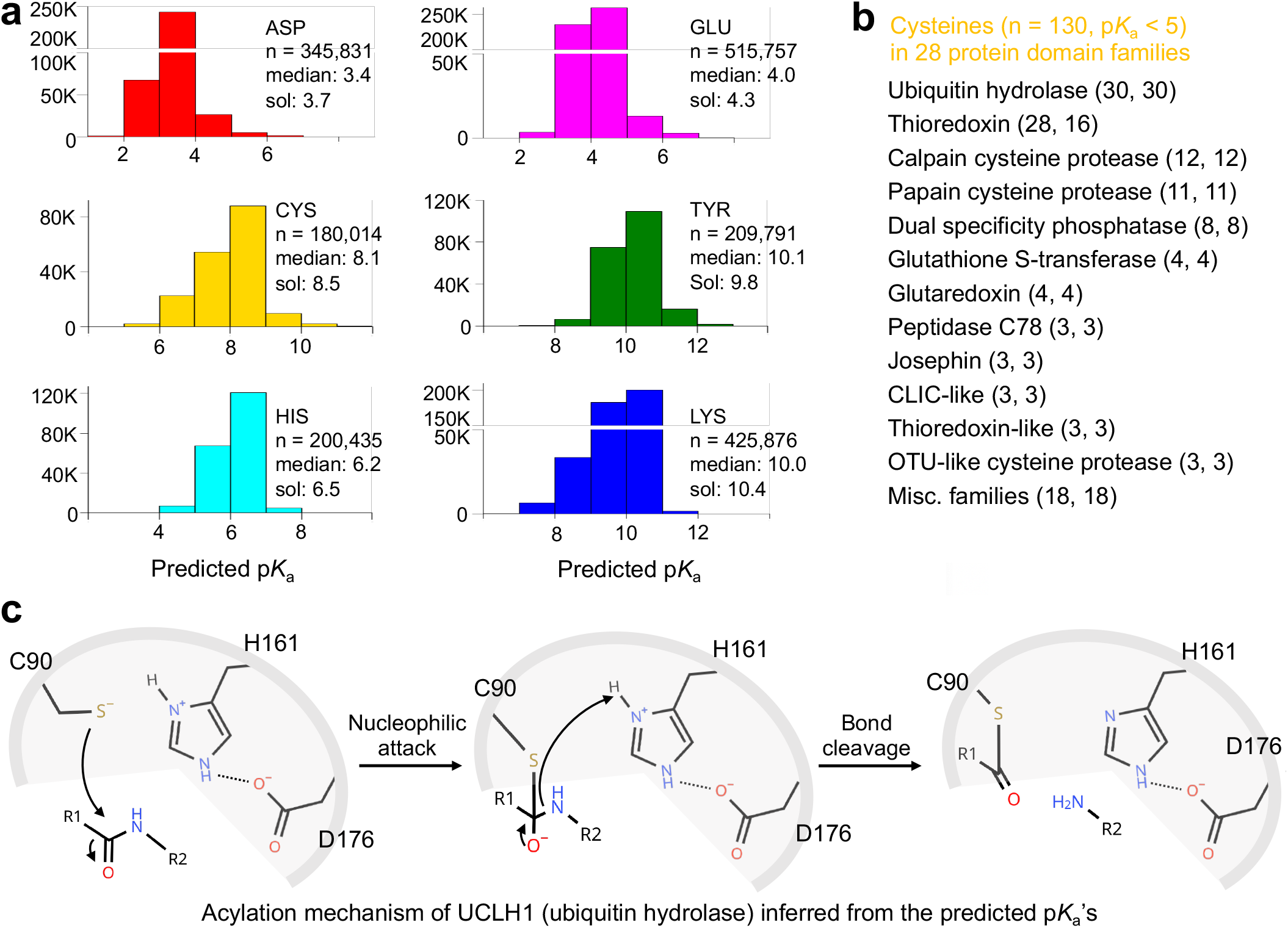
Human proteome p*K*_a_ predictions reveal functional residues and infer catalytic mechanisms. **a.** Histograms of the KaML-ESM2 predicted p*K*_a_’s of Asp, Glu, His, Cys, Tyr, and Lys residues in the human proteome. Sol: Experimental p*K*_a_’s of model tripeptides (GXG, X= titratable). ^35^ n: total number of residues. A total of 1,877,704 p*K*_a_’s of 18,192 unique human proteins (from UniProt ^36^) are shown. 1,882 proteins with sequence length ≥1022 are excluded. **b**. Pfam ^25^ analysis of cysteines (n=130) with a predicted p*K*_a_ *<*5. Domain families names are followed by the number of low-p*K*_a_ cysteines and the number of proteins. Details are given in Supplemental Data Sheet 9. **c**. Predicted p*K*_a_’s (Fig. 5) of the catalytic triad in UCLH1 support a widely accepted mechanism for cysteine proteases.

#### Functional annotations and mechanistic interpretations

To test if the p*K*_a_ predictions by KaML-ESM2 can be used to understand biology, we mapped 130 cysteines with a predicted p*K*_a_ below 5 (i.e., deprotonated at physiological pH) to protein domain families using Pfam^25^ (Fig. 6b). Remarkably, except for proteins in the thioredoxin family (many of which have two nucleophilic cysteines), there is a one-to-one mapping between cysteine and protein, which led us to test a hypothesis that these cysteines with extremely low p*K*_a_’s are functional.

We examined the low-p*K*_a_ cysteines in 30 ubiquitin hydrolase proteins, which belong to the largest domain family here. According to UniProt^36^ records, 29 of them are indeed catalytic nucleophiles, while the remaining cysteine belongs to the inactive ubiquitin hydrolase U17L4. To further investigate these cysteines, we zoomed-in on UCHL1_C90, which has a predicted p*K*_a_ of 4.7 (Fig. 5). Consistent with this prediction, a combined biochemical and crystallography study identified Cys90 as the nucleophile in a catalytic triad composed of Cys90-His161-Asp176.^38^ The predicted p*K*_a_’s of Cys90 (4.7), His161 (7.0), and Asp176 (2.4) suggest that Cys90 acts as a nucleophile and His161 as a general acid/base stabilized by Asp176, supporting the following mechanism: Cys90(-) performs a nucleophilic attack on the carbonyl of the scissile bond, forming a tetrahedral intermediate; subsequent proton transfer from His161(+) leads to bond cleavage and product release (Fig. 6c). This inferred mechanism is consistent with the X-ray structures of the active enzyme^38^ and general knowledge of cysteine proteases, illustrating the utility of KaML-ESM for functional annotation and mechanistic interpretation of proteins.

## Discussion

We trained p*K*_a_ prediction task heads on ESM2^11^ and ESMC^13^ using the GAINES-augmented PKAD-3r database. The sequence-based KaML-ESM2 and KaML-ESMC models achieved RMSE values of about 0.5 units for six titratable residue types across four benchmark test sets composed of native proteins. This performance significantly exceeds that of current structure-based models based on PB solvers, empirical approaches, and ML. A caveat of our model comparison is that alternative ML models trained on experimental data were not retrained on the same training set, and overlap between their training and test data was not verified. We conducted several analyses to understand the contributions to the strong performance of the KaML-ESM2/ESMC models. Our results showed that a shallow MLP task head is sufficient to extract p*K*_a_ values from residue embeddings, despite containing only weak p*K*_a_ signals, suggesting that the pretrained ESM2/ESMC representations play a major role in KaML’s performance. We acknowledge that it is challenging to disentangle the contribution from simple sequence patterns. Rives and coworkers demonstrated that low-resolution residue–residue contact maps emerge in ESM’s representations through self-supervised training on sequences evolved over millions of years.^11^ This correlates with the steady decrease in p*K*_a_ prediction errors as representations are extracted from deeper layers of the ESM model, suggesting that emergent structural patterns inform electrostatic features, consistent with the established understanding that p*K*_a_ values are determined by the local structural environment.^1^ Comparing model performance for naturally occurring vs. engineered OBTRUDEs, which lack evolutionary support, provides additional insight. Both KaML-ESM2/ESMC and zero-shot predictions for the engineered OBTRUDEs gave substantially larger errors than for natural residues. Taken together, these data led us to hypothesize that a protein’s sequence encodes – through its evolutionary context – not only its structure and function but also, indirectly, its electrostatic characteristics.

Another key contributor to the strong model performance is the GAINES-enhanced training strategy. Although we presented an ablation study and demonstrated the advantage of GAINES for both ESM-based and structure-based p*K*_a_ models, the validity and limitation of the label transfer strategy warrant further investigation. Keeping this in mind, the GAINES strategy may serve as a broadly applicable framework for addressing data scarcity in ML tasks involving protein-related questions, particularly given the growing representation capacities of pLMs.

ESMC was trained on 2.8 billion sequences,^13^ roughly a 40-fold increase over ESM2.^11^ Aligning with its ability to better infer emergent structures,^13^ ESMC achieved 0.2-unit lower RMSE in zero-shot predictions on holdout sets, and the trained KaML-ESMC model improved prediction of SNase OBTRUDEs p*K*_a_’s by about 0.8 units in RMSE. We demonstrated that the prediction errors for OBTRUDEs are sharply decreased when train set includes OBTRUDEs with local sequences differing significantly from the test set. This indicates that supervised learning can at least partially compensate for the lack of evolutionary information. Our benchmark analysis revealed that, for the OBTRUDE dataset, all evaluated models perform worse than the first-generation implicit-solvent CpHMD method,^34^ suggesting that later CpHMD methods such as the hybrid-solvent^39^ or all-atom CpHMD^40^ approaches would perform even better, as they more accurately describe ionization-induced local unfolding due to engineered mutations.^41^ Thus, a possible future direction is to compile a synthetic dataset of CpHMD-derived p*K*_a_ values for deeply buried residues. A related strategy, though not explicitly aimed at deeply buried residues, has been employed in the development of the DeepKa model.^7^

Sensitivity to structural perturbations is a well-known weakness of structure-based p*K*_a_ prediction methods.^1^ Since p*K*_a_ values are macroscopic properties reflecting the equilibrium of a conformational ensemble, removing the requirement for structural input is a major strength of KaML-ESMs. However, it also represents a limitation, as some proteins function by switching between two conformational states with macroscopically distinct electrostatic properties. KaML-CBT2, which is implemented in the KaML platform to predict conformation-dependent p*K*_a_ values, may benefit from incorporating sequence information or alternative architectures to improve performance. Incorporating structural information could also enable prediction of proton tautomers, which currently requires CpHMD simulations.^3,32^

We demonstrated that the KaML platform enables electrostatic characterization across the human proteome and illustrated a potential application to identify protein functional sites and deduce catalytic mechanisms. We envision integrating KaML p*K*_a_ predictions into constant-pH MD simulations^3^ to model the dynamic interplay between protonation state changes and conformational transitions critical to biological function. Such an approach would deepen our understanding of how electrostatic remodeling drives protein function — for example, in proton-coupled gating of ion channels and activation of membrane transporters.

## Methods

### Dataset and model training

KaML-ESM models were trained on the updated PKAD-3r dataset, which extends PKAD-3^8^ by incorporating 38 additional experimental p*K*_a_ values of SNase OBTRUDEs (Table 3 and Suppl. Table S1). The PKAD-3r dataset was randomly split into training and holdout test sets in a 9:1 ratio using the ‘StratifiedGroupKFold’ strategy from Scikit-learn,^18^ where each group corresponds to a unique residue (defined by a unique UniProt ID and a Uniprot sequence ID) and stratification was performed on experimental p*K*_a_ values. The training set was split into training and validation subsets in a 9:1 ratio. Stratified bins were manually verified and merged where necessary to ensure sufficient population for data partitioning. This procedure was repeated 50 times to generate 50 stratified random holdout sets, which were manually inspected to exclude any protein overlapping with the training data despite carrying a distinct UniProt ID. KaML-ESM is a MLP task head with three hidden layers built on top of the corresponding ESM foundation model. MLP training was conducted on p*K*_a_ shifts relative to the model values (penta-peptides,^35^ Asp 3.7, Glu 4.3, His 6.5, Cys 8.5, Tyr 9.8, and Lys 10.4). Pretraining used a synthetic p*K*_a_ dataset generated with our previous KaML-CBtree model.^8^ Two separate KaML-ESM models were trained for acidic (Asp, Glu, Cys, and Tyr) and basic residues (His and Lys), following our previous work.^8^

For each training-test split, 10 models were trained on independent training–validation splits in a 9:1 ratio. For each holdout test, predictions from 10 models were averaged. For the production model, 50 train–validation splits were generated from the combined PKAD-3r and external test dataset, with 10 models trained per split, resulting in a final ensemble of 500 models. More details on model architecture and training are given in Suppl. Methods.

### Ablation experiments

#### Model pretraining (PT) and separation of acid/base models (AB)

We tested four KaML-ESM2 models: (1) no PT and no AB (baseline model); (2) PT only; (3) AB only; and (4) PT and AB. Compared to the baseline model, applying PT decreased the holdout test root-mean-square error (RMSE) from 0.93 to 0.89, and applying AB decreased the test RMSE from 0.93 to 0.76 (Suppl. Table S2). Moreover, the combination of PT and AB demonstrates a synergistic effect, reducing RMSE from 0.93 to 0.73 (Suppl. Table S2).

#### Analysis of ESM layers

An earlier study found that different aspects of protein evolutionary features such as structures and functions are learned across different pLLM layers, with deeper layers attending to residue contact relationships.^21^ Considering that protein p*K*_a_’s are determined by local environment,^8^ we asked if there is a particular ESM2 layer that offers the most accurate representation of protein ionization states. To test this, we trained dedicated KaML-ESM models using embeddings from different layers of ESM2_650M, ESM2_15B, or ESMC_6B. Interestingly, the overall RMSEs of ESM2_650M and ESM2_15B models exhibit multiple local minima, while the RMSE of ESMC decreases monotonically with layer depth (Fig. S7). This difference reflects learning saturation in ESM2 models, where representation learning plateaus for several amino acids prior to the final layer (Suppl. Tables S3-S4), while ESMC maintains continuous improvement across all amino acids through the final layers (Suppl. Table S5). It is noteworthy despite its smaller architecture (33 layers and 640 M parameters), ESM2_650 M achieves comparable performance in learning protein p*K*_a_ values.

#### Contributions from ESM representations and task-specific learning

To separate the effects of the pretrained foundation model from those of task-specific training, we conducted five ablation experiments with errors evaluated on the 50 stratified random holdout sets (Suppl. Table S8). Comparing with the base line model RMSE 0.75 (KaML-ESM2 trained without pretraining and GAINES): (1) Zero-shot predictions of ESM2 gave RMSE=1.59. (2) The KaML-ESM2 with a single epoch training gave RMSE=0.90. (3) KaML-ESM2 trained using the first layer of ESM2 gave RMSE=1.22. (4) KaML-ESM2 trained with randomly shuffled p*K*_a_ labels gave RMSE=1.37. (5) An MLP task head on top of the pLLM ProstT5^22^ gave RMSE=0.89.

#### Contribution from GAINES augmentation

To verify that the performance gains from GAINES augmentation is meaningful, we conducted ablation experiments by retraining KaML-ESM2 and testing on the 50 holdout sets (Fig. 3): (1) duplicating the training set to expand its size by a factor of 10; (2) varying the representation similarity level: [0.2, 0.4], [0.4, 0.6], or [0.6, 0.8].

### Model benchmarking against alternative approaches

We benchmarked KaML-ESMs against alternative approaches using five distinct test sets (Table 3): 50 stratified random holdout sets, 20 CD-HIT-partitioned holdout sets, the EXP67S test set, the external dataset (published after April 2023), and the OBTRUDE dataset. The OBTRUDE dataset contains abnormally shifted p*K*_a_’s of 89 engineered Asp, Glu, and Lys mutations in SNase variants,^14–16^ which have previously served as a blind benchmark for p*K*_a_ prediction methods.^1^ For evaluation on EXP67S, external, and OBTRUDE datasets, KaML-ESMs were retrained after excluding overlapping residues from the training set. As alternative ML models could not be retrained, potential overlap between their training data and the test sets was not assessed unless otherwise noted.

Alternative approaches include the null model, KaML-CBT/CBT2,^8^ DeepKa,^7^ pkaani,^9^ aLCnet,^10^ PKAI/PKAI+,^5^ pKAML,^27^ ME-pKa,^42^ the PB solver Pypka,^2^ and the empirical PROPKA3 method.^4^ For the structure-based methods, the PDB files archived in the PKAD-3 database were used. All the PDB files were preprocessed to keep only the monomer containing the residue of interest. For the engineered mutants without available PDB files, structures are generated using SWISS-MODEL^43^ based on the wild type template. For sequence-based models, the input sequences were thsoe in the corresponding PDB files.

## Acknowledgment

Financial support by the National Institutes of Health (R35GM148261 and R01CA256557) is acknowledged. We thank Marius Wiggert (EvolutionaryScale, PBC, New York, NY, USA.) for advice and facilitating the usage of ESM3 and ESMC. This work was supported by an EvolutionaryScale compute grant.

## Supplemental Materials

Supplemental Materials contains Supplemental Methods, Tables, and Figures. Supplemental Data File contains 9 Data Sheets.

## Data Availability

The PKAD-3_v2 database is freely searchable and downloadable at https://database.computchem.org/pkad-3. All training and benchmark datasets are freely downloadable at https://github.com/JanaShenLab/KaML-ESM/datasets. Model weights for KaML-ESM2_650M are accessible at https://doi.org/10.5281/zenodo.17943825. Model weights for KaML-ESMC_6B are accessible at https://doi.org/10.5281/zenodo.17943447. Model weights for KaML-CBT2 are accessible at: https://doi.org/10.5281/zenodo.17943947.

## Code Availability

All KaML models are open-source and freely available for non-commercial use. The source code, pretrained model weights, documentation, and instructions are available at https://github.com/JanaShenLab/KaML-ESM.

