## Supplementary Information for "Predicting Protein Electrostatics with Protein Language Models"

**Supplementary Tables** 14

**Supplementary Figures** 22

### Supplementary Methods

#### 1. Pretraining dataset

We used KaML-CBTree[1] to predict the p$K_a$ values of all proteins in the pkPDB[2] database which are composed of proteins from the PDB. If necessary, missing structure elements were completed using PDBFixer of OpenMM.[3] In total, 5,616,944 p$K_a$ values were predicted for 1,570,916 unique residues in 32,418 unique proteins. To reduce computational cost, we created a subset as the model pretraining dataset. For each titratable amino acid, we randomly sampled predicted p$K_a$ values at a ratio of 20:1 relative to the number of entries in PKAD-3. In total, the pretraining set contains 35,080 p$K_a$'s from 10,400 Asp, 11,580 Glu, 5,860 His, 2,503 Cys, 820 Tyr, and 3,917 Lys in 9,945 proteins. Residues in the PKAD-3 database were excluded from the pretraining dataset to avoid data leakage in later model training/testing.

To ensure a precise correspondence between the sequence and the protein studied by experiment regardless of the residue-specific resolution, we extracted sequences directly from the PDB files. This extraction process involved identifying resolved residues from ATOM records while detecting unresolved segments through multiple methods: analyzing discontinuities in residue numbering, identifying unusually large distances between adjacent backbone atoms, and incorporating segments explicitly annotated as missing in the

PDB metadata (`REMARK 465`). Non-standard amino acids were carefully mapped to their closest canonical equivalents (e.g., selenomethionine [MSE] to methionine [M], methyllysine [MLY] to lysine [K], phosphoserine [SEP] to serine [S]), or otherwise represented as `X` when no direct canonical mapping exists. This rigorous and automated sequence extraction procedure was implemented using the `extract_sequence.py` script from *PDB doctor*, an unpublished in-house software suite available at https://github.com/wayyne/pdb_doctor.

#### 2. Training and benchmark datasets

**The PKAD-3r database.** We recently published the PKAD-3 database,[1] which significantly expands upon the PKAD-2 database[4] recently developed by the Alexov lab (Table S1). The PKAD-3 database includes 1,167 p$K_a$'s of 992 unique residues in 247 proteins (wild type or mutant).[1] In this work, we extended the PKAD-3 database by including additional 38 p$K_a$ values of 38 engineered OBTRUDEs of staphylococcal nuclease (SNase) missed in the previous literature search.[5,6] The revised database PKAD-3r (Table S1) includes 1,205 p$K_a$'s of 1030 unique residues in 285 proteins (wild type or mutant). This dataset serves as a foundation for model training, benchmarking, and data augmentation.

**Training and holdout data partitions.** We used the same data partitioning protocol as in the development of KaML-CBTree and KaML-GAT models.[1] Briefly, we labeled a unique residue as a unique combination of Uniprot ID + the Uniprot residue ID (Uni_resid). To account for mutants and multiple conformational states, we defined these proteins as 'Uniprot ID-mutation or conformational state'. For example, P0AEG4-H32L represents H32L mutation on DsbA (P0AEG4). For SNase (P00644), which has multiple background constructs, e.g., WT, PHS, and Δ+PHS, we used P00644-V66D, P00644-WT-V66D, and P00644-PHS-V66D to represent the Δ+PHS, WT, and PHS constructs, respectively.

Data partitioning into training, validation, and holdout test sets used the Stratified-

GroupKFold strategy from the scikit-learn Python package.[7] Here, a group corresponds to a unique residue, and stratification was based on experimental p$K_a$ values. Stratified bins were manually verified and merged where necessary to ensure sufficient population for train-validation-test partitioning. From the PKAD-3r dataset,[1] we first randomly sampled 10% as the holdout test set with the 'StratifiedGroupKFold' strategy. The remaining 90% was split into training and validation sets in a 9:1 ratio using the same protocol, repeated 20 or 50 times independently. The same 9:1 split was applied to the pretraining dataset.

**Local sequence similarity calculation.** Local sequence similarity based on a 21 residue-window centered on the residue of interest was calculated to check for potential data leakage between training/validation and test sets. Sequences are aligned using the Biopython PairwiseAligner with BLOSUM62 substitution matrix.

**A new external test set.** Note, the PKAD-3r database only includes experimental p$K_a$ values published before April 2023. To compile an external evaluation dataset, we searched PubMed for publications after April 2023 using 'p$K_a$' as a title/abstract keyword. We then manually verified the reported p$K_a$ values. We also identified NMR-derived p$K_a$ values for histidines in SNase mutants[8] that were missed during PKAD-3 curation. When measurements at multiple KCl concentrations were available, the value at 0.1 M was recorded. In total, 54 p$K_a$ values (39 His, 1 Cys, 1 Tyr, and 13 Lys) from 17 proteins were compiled as an external test set. These data were excluded from model training and GAINES augmentation. In total, 54 p$K_a$'s (39 His, 1 Cys, 1 Tyr, and 13 Lys) from 17 proteins were compiled and used exclusively as an external test set, withheld from model training and GAINES augmentation.

**The SNase OBTRUDE dataset.** The SNase OBTRUDE dataset contains p$K_a$'s derived from NMR shifts or folding stability changes for 89 titratable substitutions of buried hydrophobic residues in hyperstabilized SNase variants.[5,6] Most OBTRUDE residues show very large (over 2 pH units) p$K_a$ shifts relative to the model p$K_a$'s, while a few lysines show no p$K_a$ shifts.

**A new searchable database PKAD-3_v2 for the community.** The external test set was combined with PKAD-3r and released as PKAD-3_v2 on the website (http://database.computchem.org/pkad-3). In total, the PKAD-3_v2 database contains 1,259 p$K_a$'s of 1084 unique residues in 302 wild type or mutant proteins. Currently, only the first 50 entries are displayed, but the entire database is searchable and downloadable.

#### 3. The GAINES protocol

To address the challenge of limited training data for ML, we introduced GAINES (auGment dAta wIth lateNt spacE Sampling), an approach that enriches model training by augmenting the dataset with synthetic labels assigned via a biophysically motivated sampling strategy. Specifically, GAINES leverages token embeddings of query residues from the original training set to locate value residues in an unrelated database that are similar in the latent space and often dissimilar at the sequence level. We computed per-residue embeddings (see section Model architecture) for every protein archived in the RCSB Protein Data Bank and then calculated cosine similarity between their residues and the query residues in the PKAD-3r database. Any residue with a similarity score greater than 0.8 is designated as a value residue, assigned the label (e.g., p$K_a$) of the corresponding query residue, and added to the training dataset. This method is motivated by the observation that residue-level embeddings encode local environments, which govern protein p$K_a$ values. Importantly, synthetic data points are used exclusively during training and not included in the test set.

#### 4. Model architecture

The KaML-ESM models were built and trained using the PyTorch package.[9] A multilayer perceptron (MLP) with 3 hidden layers was built as a task head on top of an ESM model. The number of neurons in KaML-ESM2_650M, KaML-ESM2_15B, and KaML_ESMC models was 1280-512-256-32-1, 5120-2048-1024-256-1, and 2560-1024-512-64-1, respectively. The rectified linear unit (ReLU) activation function is applied to each hidden layer to add non-linearity. The output of the last neuron is converted to the prediction through a linear activation function. Batch normalization is applied to all layers except the last one. For model training, we used a learning rate of 0.0005 and a dropout rate of 0.2 with a batch size of 64 and a maximum epoch of 200. The Adam optimizer is used to minimize the mean squared error (MSE) loss. Early stopping is applied to prevent overfitting: model training is terminated if the MSE loss does not decrease within the next 10 epochs with a tolerance of 0.1.

The evolutionary scale models (ESMs) are protein large language models (pLLMs) based on the BERT[10] style transformer[11] architecture. We used ESM2[12] and ESMC,[13] which are encoder-only transformers trained using masked learning of 50 million protein sequences. ESM2_t33_650M_UR50D (ESM_650M) model has 33 transformer layers (20 attention heads each) and 650 million parameters. ESM2_t48_15B_UR50D (ESM2_15B) has 48 transformer layers (40 attention heads each) and 15 billion parameters. ESMC-6B-2024-12 (ESMC) has 80 transformer layers (40 attention heads each) and 6 billion parameters. This model, which is focused on representation learning of the underlying biology, is a parallel model to the generative model ESM3.[14] For titratable residues of interest, the token embeddings were extracted from ESM2_650M, ESM2_15B, or ESMC, which have dimensions of 1280, 5120, and 2560, respectively. Note, ESM2_650M can be loaded to GPUs with $\sim 10$ GB memory while the ESM2-15B requires a memory space of $\sim 100$ GB and ESMC embeddings can only be extracted using the forge-API (https://forge.evolutionaryscale.ai/) with daily token and speed limitations.

**Parameter-efficient fine-tuning of ESM2_650M.** We tested whether parameter-efficient fine-tuning of ESM2_650M improved task head performance relative to training with frozen embeddings. Fine-tuning was performed using the qLoRA method[15] on the `ESM2_t33_650M_UR50D` backbone. LoRA adapters were applied throughout the ESM2 transformer using rank $r = 64$, scaling factor $\alpha = 8$, and adapter dropout of $0.1$. A quantization block size of 32, bfloat16 compute precision, double quantization, gradient checkpointing, and a paged 8-bit AdamW optimizer were used.

The same KaML-ESM2_650M MLP task head architecture described above was used for the fine-tuning experiments. The task was formulated as residue-level regression and optimized using mean squared error loss. Training used a batch size of 4, a learning rate of $5 \times 10^{-5}$ for both the qLoRA adapter parameters and the task head, and weight decay of $1 \times 10^{-4}$. Models were trained for 50 total epochs. During the first 10 epochs, the qLoRA adapters were disabled and only the task head was optimized. During the remaining epochs, the qLoRA adapters and task head were optimized jointly. The pretrained ESM2 backbone weights were not directly updated. The final model was selected by validation loss, without early stopping.

#### 5. Model training protocol

Due to the distinct physiochemical properties of acidic (Asp, Glu, Cys, and Tyr) and basic (His and Lys) residues, we trained acid and base models separately. This strategy has been shown to enhance model performance.[1] The models were trained on the p$K_a$ shifts relative to the solution p$K_a$ values of penta-peptides $CH_3COAAXAANH_2$.[16] Accordingly, the solution p$K_a$'s are 3.7 for Asp, 4.3 for Glu, 6.5 for His, 8.5 for Cys, 9.8 for Tyr, and 10.4 for Lys. The p$K_a$ shifts are converted back to p$K_a$ values after the model training or prediction. We pretrained our model using a synthetic dataset composed of p$K_a$ values predicted by the structure-based KaML-CBtree model.[1] The Glorot algorithm[17] was used to initialize the model weights in the pretraining stage. For each training/validation-holdout split, 10 models with were trained on independent 9:1 training-validation splits. Each prediction is thus the average of an ensemble of 10 models. In the production stage, we retrained KaML-ESM2/ESMC models using the entire PKAD-3_v2 dataset (PKAD-3r + external set) for training and CV. With 50 training-validation splits and 10 models per split, the final model is an ensemble of 500 models.

#### 6. Implementation of the KaML platform

The KaML-ESM platform comprises multiple modular stages managed within a unified Python framework, facilitating ease of use and reproducibility. Protein sequences or structures (specified via UniProt identifiers, PDB files, PDB IDs, or FASTA sequences) are initially retrieved through established web services (UniProt, RCSB) or processed directly when provided by users. For sequences without structural information, computational folding is performed using the remote ESM3-medium model accessible through the Forge service.

Following retrieval and preprocessing, sequences and structures are parsed to identify titratable residues. Embeddings are subsequently computed from these sequences using a pLLM, specifically ESM2 (esm2_t33_650M_UR50D)[12] or ESMC (esmc-6b-2024-12),[13] with model selection determined by residue type (acidic or basic). By default, embeddings for acidic residues are extracted from ESM2 and processed using a multi-layer perceptron (MLP) configured as 1280→512→256→32→1 neurons with dropout regularization. Embeddings for basic residues are derived from ESMC and processed through a separate MLP architecture of 2560→1024→512→64→1 neurons, similarly utilizing dropout regularization. Details are given in paragraph Architecture of the KaML-ESM models.

Predictions from acidic and basic channels are generated via parallelized inference using ensemble models, supported by statistical normalization strategies developed from extensive training datasets. The acidic and basic channel predictions are consolidated to yield the predicted p$K_a$ values, shifts, and standard error metrics. An additional, independently operated modular conformer channel is integrated, allowing users to employ alternative conformer-sensitive predictive models in cases where distinct conformations significantly impact predicted p$K_a$ values. Currently, this channel employs the previously validated KaMLs-CBTree model.[1] Predictions from the conformer channel are reported independently in the final output.

**The command-line KaML platform.** The KaML platform (https://github.com/JanaShenLab/KaML-ESM) is openly available, accompanied by pretrained MLP ensemble weights. Specifically provided are pretrained weights for the acidic ESMC channel and for cysteine residues using layer 33 of the ESM2 model. The platform is designed to offer users flexibility in specifying foundational models independently for each prediction channel. For example, users can override default settings to utilize ESMC for both acidic and basic channels when analyzing sequences exceeding the 1022-residue limit of ESM2, as ESMC supports sequence lengths up to 2046 residues. In these scenarios, the platform automatically utilizes ESMC for the acidic channel, emphasizing the practical benefit of including these pretrained weights. Additionally, users may optionally select alternative layers within ESM2, such as employing layer 33 specifically for cysteine residues, reflecting enhanced performance demonstrated in the primary analysis. By default, however, the platform utilizes layer 31.

Finally, predictions are integrated back into structural outputs by annotating the structure files with residue-specific predictions in the B-factor field, facilitating visualization and downstream structural analysis. Prediction results, distinguishing between acidic/basic channel outputs and independent conformer channel predictions, are documented in CSV files, enhancing clarity and accessibility for further analysis.

**The browser-based KaML application.** We provide an easy-to-use online browser-based GUI (https://kaml.computchem.org/) for non-commercial academic and research purposes. Users can provide either the protein sequence, Uniprot ID, PDB ID, or PDB files of interest. By clicking the 'Run pipeline' button, the complete KaML-ESM model will run on our web server. A PDB format file with the B-factor column filled with the predicted p$K_a$ shifts and a CSV file with predicted p$K_a$ values for all titratable residues can be downloaded. The screenshots of the web KaML application is given in Suppl. Fig. S14.

#### 7. Prediction of p$K_a$ values for the human proteome

We applied KaML-ESM2 to predict the p$K_a$ values of Asp, Glu, Cys, Tyr, His, and Lys side chains for 18,192 (out of a total of 20,074) human proteins in UniProt with sequence lengths $\leq$ 1,022 amino acids (the sequence length limit for ESM2). In total, the p$K_a$ values of 345,831 Asp, 515,757 Glu, 180,013 Cys, 209,791 Tyr, 200,435 His, and 425,876 Lys were recorded.

#### 8. Ablation studies

To evaluate the contributions from pre-training (PT) and separate acid/base (AB) training, we tested four KaML-ESM2 models: (1) no PT and no AB (baseline model); (2) PT only; (3) AB only; and (4) PT and AB. The combination of PT and AB demonstrated a synergistic effect (Suppl. Table S2).

To disentangle contributions from the pretrained foundation model from those of task-specific training, we carried out five ablation experiments using ESM2 and 50 independent holdout test sets (Suppl. Table S8). (1) To demonstrate the performance of the foundation model itself, zero-shot predictions were generated without performing any MLP training. (2) To demonstrate the effect of task-specific training, we trained an MLP for a single epoch. (3) To demonstrate the impact of the foundation model's learned representations, an MLP was trained using the first layer of ESM2. (4) To verify that the training task itself is important, an MLP was trained on the same ESM2 embeddings but with the p$K_a$ labels randomly permuted. (5) To compare with alternative foundation models, an MLP leveraging the embeddings of the recent protein LLM ProstT5[18] were trained using the same settings.

To verify that the performance gains from GAINES augmentation is meaningful rather than merely from increased data volume, we conducted three ablation experiments (Suppl. Table S2): (1) duplicating the training set to expand its size by a factor of 10; (2) adjusting the similarity threshold to the ranges [0.2, 0.4], [0.4, 0.6], and [0.6, 0.8], and comparing retrained models against the baseline model using the similarity threshold of $>$0.8; (3) comparing the RMSE of residues with GAINES-augmented data to those without such augmentation.

#### 9. Model performance comparison against alternative approaches

We evaluated KaML-ESMs on five different datasets: 50 stratified holdout sets, 20 CD-HIT-partitioned hold sets, the EXP67S dataset, an external test set (published after April 2023), and the SNase OBTRUDE test set. These datasets are available at `https://github.com/JanaShenLab/KaML-ESM/tree/main`. Alternative approaches include NULL model, DeepKa,[19] pkaani,[20] aLCnet,[21] PKAI/PKAI+,[22] pKAML,[23] and ME-pKa,[24] physics-based Pypka,[25] and empirical PROPKA3 method.[26] For structure-based methods, the PDB files archived in the PKAD-3 database are used. All the PDB files are preprocessed to keep only the monomer containing the residue of interest. For the engineered mutants without available PDB files, structures are generated using SWISS-MODEL[27] based on the wild type template. For sequence-dependent methods, the input sequences are generated based on the corresponding PDB files.

Predictions of pKAML[23] were generated using the pKAML web server (`https://onodalab.ees.hokudai.ac.jp/pkalm`) Predictions of DeepKa[19] were generated using the DeepKa web server (`http://www.computbiophys.com/DeepKa/main`). Predictions of pkaani[20] wre generated using the command-line version downloaded from Github (`https://github.com/isayevlab/pKa-ANI`). Predictions of aLCnet were generated using the command-line version downloaded from Github (https://github.com/feiglab/ProteinStructureEmbedding). Predictions of PKAI and PKAI+ are generated using the command-line version downloaded from Github (https://github.com/bayer-science-for-a-better-life/pKAI). Predictions of Pypka were generated using the command-line version downloaded from Github (https://github.com/mms-fcul/PypKa). Predictions of PROPKA3 were generated using the command-line version downloaded from Github (https://github.com/jensengroup/propka). Note, the model performances from Me-pKa are author-reported with different data splitting schemes.

### Supplementary Tables

Table S1: Summary of p$K_a$ datasets and usage in this work

| Dataset | Proteins | Residues | p$K_a$'s | Usage |
|---|---|---|---|---|
| Experimental data | | | | |
| PKAD-3r | 285 | 1,030 | 1,205 | Training KaML-ESMs |
| PKAD-3r w/o OBTRUDE | 205 | 941 | 1,116 | Training KaML-ESMs for testing on OBTRUDE |
| OBTRUDE | 89 | 89 | 89 | Testing |
| External test set | 17 | 54 | 54 | Testing |
| PKAD-3r+External | 302 | 1,084 | 1,259 | Training production KaML-ESMs |
| Synthetic data | | | | |
| KaML-CBT predicted p$K_a$'s | 9,945 | 29,457 | 35,080 | Pretraining |
| GAINES data | 2,164 | 4,220 | 4,220 | Augmentation |

The number of unique proteins, residues, and p$K_a$'s are given here. The OBTRUDE and external validation datasets are given in Supplemental Data file. The online database is revised to include PKAD-3r and the external validation dataset, which together are referred to as PKAD-3_v2.

Table S2: Ablation study of pretraining and acid-base model separation for KaML-ESM2 p$K_a$ predictions

| | All | Asp | Glu | Cys | Tyr | His | Lys |
|---|---|---|---|---|---|---|---|
| **PT + AB** | $0.73 \pm 0.03$ | $0.72 \pm 0.06$ | $0.67 \pm 0.04$ | $1.01 \pm 0.09$ | $1.65 \pm 0.17$ | $0.67 \pm 0.03$ | $0.55 \pm 0.04$ |
| No PT + No AB | $0.93 \pm 0.04$ | $0.95 \pm 0.09$ | $0.71 \pm 0.04$ | $1.03 \pm 0.11$ | $1.54 \pm 0.19$ | $0.96 \pm 0.05$ | $0.99 \pm 0.09$ |
| PT only | $0.89 \pm 0.03$ | $0.93 \pm 0.08$ | $0.73 \pm 0.04$ | $1.05 \pm 0.11$ | $1.44 \pm 0.17$ | $0.82 \pm 0.04$ | $0.88 \pm 0.10$ |
| AB only | $0.76 \pm 0.02$ | $0.71 \pm 0.05$ | $0.70 \pm 0.04$ | $1.00 \pm 0.09$ | $1.74 \pm 0.19$ | $0.79 \pm 0.12$ | $0.59 \pm 0.05$ |

Overall and amino acid-specific RMSE±standard error of the p$K_a$ predictions by the KaML-ESM2 model using 20 hold-out tests (same as in our previous work[1]). PT: pretraining, AB: training acidic and basic models separately. Residue embeddings were extracted from ESM2_650M layer 33. Note: no GAINES augmentation was used.

Table S3: p$K_a$ prediction errors of the models trained with the embeddings from different layers of ESM2_650M

| | All | Asp | Glu | Cys | Tyr | His | Lys |
|---|---|---|---|---|---|---|---|
| | | | | Without pretraining | | | |
| layer33 | $0.76 \pm 0.02$ | $0.71 \pm 0.05$ | $0.70 \pm 0.04$ | $1.00 \pm 0.09$ | $1.74 \pm 0.19$ | $0.79 \pm 0.04$ | $0.59 \pm 0.05$ |
| layer32 | $0.77 \pm 0.02$ | $0.70 \pm 0.04$ | $0.69 \pm 0.04$ | $1.04 \pm 0.09$ | $1.75 \pm 0.20$ | $0.84 \pm 0.04$ | $0.61 \pm 0.05$ |
| **layer31** | $0.75 \pm 0.02$ | $0.68 \pm 0.04$ | $0.62 \pm 0.03$ | $1.23 \pm 0.11$ | $1.58 \pm 0.17$ | $0.81 \pm 0.05$ | $0.71 \pm 0.07$ |
| layer30 | $0.79 \pm 0.02$ | $0.71 \pm 0.04$ | $0.74 \pm 0.03$ | $1.15 \pm 0.10$ | $1.43 \pm 0.18$ | $0.85 \pm 0.04$ | $0.65 \pm 0.05$ |
| layer29 | $0.82 \pm 0.02$ | $0.74 \pm 0.03$ | $0.74 \pm 0.04$ | $1.30 \pm 0.14$ | $1.50 \pm 0.20$ | $0.92 \pm 0.05$ | $0.62 \pm 0.06$ |
| layer28 | $0.81 \pm 0.02$ | $0.73 \pm 0.04$ | $0.75 \pm 0.04$ | $1.23 \pm 0.11$ | $1.49 \pm 0.22$ | $0.89 \pm 0.05$ | $0.61 \pm 0.06$ |
| layer27 | $0.81 \pm 0.02$ | $0.74 \pm 0.04$ | $0.74 \pm 0.04$ | $1.25 \pm 0.11$ | $1.56 \pm 0.22$ | $0.89 \pm 0.05$ | $0.64 \pm 0.06$ |
| layer26 | $0.86 \pm 0.03$ | $0.78 \pm 0.04$ | $0.79 \pm 0.04$ | $1.36 \pm 0.13$ | $1.66 \pm 0.25$ | $0.90 \pm 0.04$ | $0.70 \pm 0.07$ |
| layer25 | $0.87 \pm 0.03$ | $0.81 \pm 0.04$ | $0.79 \pm 0.04$ | $1.48 \pm 0.13$ | $1.71 \pm 0.26$ | $0.91 \pm 0.04$ | $0.68 \pm 0.06$ |
| layer24 | $0.87 \pm 0.03$ | $0.83 \pm 0.04$ | $0.81 \pm 0.04$ | $1.36 \pm 0.13$ | $1.58 \pm 0.24$ | $0.91 \pm 0.04$ | $0.68 \pm 0.07$ |
| layer23 | $0.89 \pm 0.03$ | $0.85 \pm 0.04$ | $0.82 \pm 0.04$ | $1.42 \pm 0.13$ | $1.57 \pm 0.23$ | $0.90 \pm 0.04$ | $0.71 \pm 0.07$ |
| layer22 | $0.89 \pm 0.03$ | $0.84 \pm 0.04$ | $0.81 \pm 0.04$ | $1.45 \pm 0.14$ | $1.49 \pm 0.23$ | $0.91 \pm 0.04$ | $0.72 \pm 0.07$ |
| layer21 | $0.87 \pm 0.02$ | $0.84 \pm 0.03$ | $0.76 \pm 0.04$ | $1.43 \pm 0.12$ | $1.49 \pm 0.20$ | $0.92 \pm 0.04$ | $0.74 \pm 0.08$ |
| layer20 | $0.89 \pm 0.03$ | $0.83 \pm 0.04$ | $0.80 \pm 0.04$ | $1.45 \pm 0.16$ | $1.64 \pm 0.21$ | $0.91 \pm 0.04$ | $0.79 \pm 0.07$ |
| layer19 | $0.88 \pm 0.03$ | $0.83 \pm 0.04$ | $0.77 \pm 0.04$ | $1.41 \pm 0.16$ | $1.76 \pm 0.25$ | $0.92 \pm 0.04$ | $0.79 \pm 0.07$ |
| layer18 | $0.87 \pm 0.03$ | $0.84 \pm 0.04$ | $0.77 \pm 0.04$ | $1.43 \pm 0.14$ | $1.60 \pm 0.22$ | $0.88 \pm 0.04$ | $0.80 \pm 0.08$ |
| layer17 | $0.87 \pm 0.03$ | $0.85 \pm 0.04$ | $0.76 \pm 0.04$ | $1.58 \pm 0.12$ | $1.51 \pm 0.22$ | $0.89 \pm 0.04$ | $0.79 \pm 0.07$ |
| | | | | With pretraining | | | |
| layer33 | $0.73 \pm 0.03$ | $0.72 \pm 0.06$ | $0.67 \pm 0.04$ | $1.01 \pm 0.09$ | $1.65 \pm 0.17$ | $0.67 \pm 0.03$ | $0.55 \pm 0.04$ |
| layer32 | $0.71 \pm 0.03$ | $0.69 \pm 0.05$ | $0.65 \pm 0.04$ | $1.14 \pm 0.10$ | $1.47 \pm 0.14$ | $0.67 \pm 0.03$ | $0.61 \pm 0.04$ |
| **layer31** | $0.70 \pm 0.02$ | $0.65 \pm 0.04$ | $0.61 \pm 0.04$ | $1.11 \pm 0.09$ | $1.18 \pm 0.16$ | $0.68 \pm 0.03$ | $0.69 \pm 0.07$ |
| layer30 | $0.75 \pm 0.02$ | $0.71 \pm 0.04$ | $0.75 \pm 0.04$ | $1.18 \pm 0.11$ | $1.34 \pm 0.16$ | $0.65 \pm 0.04$ | $0.62 \pm 0.06$ |
| layer29 | $0.75 \pm 0.02$ | $0.70 \pm 0.04$ | $0.72 \pm 0.04$ | $1.32 \pm 0.12$ | $1.35 \pm 0.18$ | $0.69 \pm 0.03$ | $0.55 \pm 0.05$ |
| layer28 | $0.74 \pm 0.02$ | $0.70 \pm 0.04$ | $0.72 \pm 0.04$ | $1.15 \pm 0.08$ | $1.39 \pm 0.19$ | $0.73 \pm 0.03$ | $0.50 \pm 0.04$ |

Overall and amino acid-specific RMSE±standard error of the p$K_a$ predictions by the KaML-ESM2 model using 20 hold-out tests (same as in our previous work[1]). Residue embeddings were extracted from the last 6 layers of ESM2_650M. Layer 31 gives the lowest overall RMSE, while the lowest RMSEs (red) for Cys, Tyr, His, and Lys p$K_a$ predictions are from other layers. Pre-training was performed for all models.

Table S4: p$K_a$ prediction errors of the models trained with the embeddings from different layers of ESM2_15B (no pretraining)

| | All | Asp | Glu | Cys | Tyr | His | Lys |
|---|---|---|---|---|---|---|---|
| layer48 | $0.74 \pm 0.03$ | $0.70 \pm 0.05$ | $0.60 \pm 0.03$ | $1.16 \pm 0.08$ | $1.70 \pm 0.20$ | $0.84 \pm 0.04$ | $0.61 \pm 0.05$ |
| **layer47** | $0.73 \pm 0.02$ | $0.68 \pm 0.04$ | $0.60 \pm 0.02$ | $1.07 \pm 0.08$ | $1.73 \pm 0.20$ | $0.85 \pm 0.04$ | $0.64 \pm 0.06$ |
| layer46 | $0.74 \pm 0.02$ | $0.69 \pm 0.04$ | $0.61 \pm 0.03$ | $1.09 \pm 0.09$ | $1.62 \pm 0.20$ | $0.85 \pm 0.04$ | $0.62 \pm 0.06$ |
| layer45 | $0.74 \pm 0.03$ | $0.70 \pm 0.04$ | $0.62 \pm 0.04$ | $1.07 \pm 0.10$ | $1.51 \pm 0.20$ | $0.85 \pm 0.04$ | $0.63 \pm 0.05$ |
| layer44 | $0.76 \pm 0.03$ | $0.68 \pm 0.04$ | $0.66 \pm 0.04$ | $1.11 \pm 0.10$ | $1.53 \pm 0.21$ | $0.87 \pm 0.04$ | $0.65 \pm 0.06$ |
| layer43 | $0.76 \pm 0.03$ | $0.70 \pm 0.04$ | $0.66 \pm 0.04$ | $1.10 \pm 0.11$ | $1.48 \pm 0.22$ | $0.87 \pm 0.04$ | $0.65 \pm 0.06$ |
| layer42 | $0.77 \pm 0.03$ | $0.70 \pm 0.04$ | $0.66 \pm 0.04$ | $1.10 \pm 0.12$ | $1.47 \pm 0.23$ | $0.90 \pm 0.04$ | $0.64 \pm 0.06$ |

Overall and amino acid-specific RMSE±standard error of the p$K_a$ predictions by the KaML-ESM2 model using 20 hold-out tests (same as in our previous work[1]). The layer that gives the lowest overall RMSE is highlighted in bold font. The lowest amino acid-specific RMSE is highlighted in red.

Table S5: p$K_a$ prediction errors of the the models trained with the embeddings from different layers of KaML-ESMC (no pretraining)

| | All | Asp | Glu | Cys | Tyr | His | Lys |
|---|---|---|---|---|---|---|---|
| **layer80** | $0.70 \pm 0.03$ | $0.64 \pm 0.04$ | $0.61 \pm 0.03$ | $1.22 \pm 0.13$ | $1.46 \pm 0.17$ | $0.74 \pm 0.04$ | $0.53 \pm 0.04$ |
| layer79 | $0.74 \pm 0.03$ | $0.68 \pm 0.05$ | $0.64 \pm 0.03$ | $1.47 \pm 0.12$ | $1.41 \pm 0.16$ | $0.74 \pm 0.04$ | $0.58 \pm 0.04$ |
| layer78 | $0.73 \pm 0.03$ | $0.67 \pm 0.05$ | $0.62 \pm 0.03$ | $1.30 \pm 0.12$ | $1.58 \pm 0.21$ | $0.77 \pm 0.04$ | $0.57 \pm 0.04$ |
| layer77 | $0.76 \pm 0.03$ | $0.69 \pm 0.05$ | $0.64 \pm 0.03$ | $1.31 \pm 0.15$ | $1.73 \pm 0.26$ | $0.79 \pm 0.04$ | $0.58 \pm 0.04$ |
| layer76 | $0.75 \pm 0.03$ | $0.67 \pm 0.05$ | $0.64 \pm 0.03$ | $1.35 \pm 0.14$ | $1.75 \pm 0.25$ | $0.76 \pm 0.04$ | $0.60 \pm 0.04$ |
| layer75 | $0.75 \pm 0.03$ | $0.67 \pm 0.05$ | $0.66 \pm 0.04$ | $1.36 \pm 0.15$ | $1.71 \pm 0.25$ | $0.78 \pm 0.04$ | $0.59 \pm 0.04$ |
| layer74 | $0.77 \pm 0.03$ | $0.69 \pm 0.05$ | $0.65 \pm 0.04$ | $1.39 \pm 0.14$ | $1.68 \pm 0.25$ | $0.83 \pm 0.04$ | $0.57 \pm 0.04$ |
| layer73 | $0.78 \pm 0.03$ | $0.70 \pm 0.06$ | $0.68 \pm 0.04$ | $1.42 \pm 0.12$ | $1.61 \pm 0.23$ | $0.84 \pm 0.04$ | $0.57 \pm 0.05$ |
| layer72 | $0.78 \pm 0.03$ | $0.70 \pm 0.06$ | $0.70 \pm 0.04$ | $1.33 \pm 0.13$ | $1.55 \pm 0.24$ | $0.84 \pm 0.03$ | $0.58 \pm 0.06$ |
| layer71 | $0.78 \pm 0.03$ | $0.70 \pm 0.05$ | $0.69 \pm 0.04$ | $1.36 \pm 0.11$ | $1.52 \pm 0.24$ | $0.83 \pm 0.03$ | $0.60 \pm 0.05$ |
| layer70 | $0.79 \pm 0.03$ | $0.70 \pm 0.05$ | $0.71 \pm 0.04$ | $1.40 \pm 0.12$ | $1.59 \pm 0.23$ | $0.83 \pm 0.04$ | $0.63 \pm 0.06$ |
| layer69 | $0.80 \pm 0.03$ | $0.72 \pm 0.06$ | $0.71 \pm 0.04$ | $1.47 \pm 0.11$ | $1.53 \pm 0.22$ | $0.84 \pm 0.03$ | $0.64 \pm 0.06$ |
| layer68 | $0.80 \pm 0.03$ | $0.72 \pm 0.06$ | $0.72 \pm 0.04$ | $1.44 \pm 0.12$ | $1.47 \pm 0.22$ | $0.84 \pm 0.04$ | $0.66 \pm 0.06$ |
| layer67 | $0.81 \pm 0.03$ | $0.73 \pm 0.06$ | $0.73 \pm 0.04$ | $1.46 \pm 0.13$ | $1.49 \pm 0.23$ | $0.85 \pm 0.04$ | $0.66 \pm 0.07$ |
| layer66 | $0.81 \pm 0.03$ | $0.74 \pm 0.05$ | $0.73 \pm 0.04$ | $1.46 \pm 0.14$ | $1.44 \pm 0.23$ | $0.85 \pm 0.03$ | $0.67 \pm 0.06$ |
| layer65 | $0.81 \pm 0.03$ | $0.73 \pm 0.05$ | $0.74 \pm 0.04$ | $1.44 \pm 0.13$ | $1.47 \pm 0.24$ | $0.85 \pm 0.04$ | $0.67 \pm 0.06$ |
| layer64 | $0.82 \pm 0.03$ | $0.75 \pm 0.06$ | $0.74 \pm 0.04$ | $1.46 \pm 0.12$ | $1.49 \pm 0.24$ | $0.86 \pm 0.04$ | $0.67 \pm 0.06$ |
| layer63 | $0.82 \pm 0.03$ | $0.75 \pm 0.06$ | $0.74 \pm 0.04$ | $1.45 \pm 0.12$ | $1.55 \pm 0.24$ | $0.85 \pm 0.04$ | $0.69 \pm 0.06$ |
| layer62 | $0.82 \pm 0.03$ | $0.76 \pm 0.06$ | $0.74 \pm 0.04$ | $1.49 \pm 0.13$ | $1.52 \pm 0.23$ | $0.85 \pm 0.04$ | $0.69 \pm 0.06$ |
| layer61 | $0.83 \pm 0.03$ | $0.77 \pm 0.06$ | $0.73 \pm 0.04$ | $1.48 \pm 0.12$ | $1.47 \pm 0.21$ | $0.86 \pm 0.04$ | $0.71 \pm 0.06$ |
| layer60 | $0.83 \pm 0.03$ | $0.76 \pm 0.06$ | $0.74 \pm 0.04$ | $1.46 \pm 0.11$ | $1.49 \pm 0.23$ | $0.87 \pm 0.04$ | $0.71 \pm 0.06$ |

Overall and amino acid-specific RMSE±standard error of the p$K_a$ predictions by the KaML-ESM2 model using 20 hold-out tests (same as in our previous work[1]). The layer that gives the lowest overall RMSE is highlighted in bold font. The lowest amino acid-specific RMSE is highlighted in red.

Table S6: Performance of the fine-tuned KaML-ESM2 model[a]

| | Overall | Asp | Glu | Cys | Tyr | His | Lys |
|---|---|---|---|---|---|---|---|
| | | | Holdouts[a] | | | | |
| Frozen | $0.55$ | $0.80$ | $0.39$ | $0.25$ | $0.54$ | $0.32$ | $0.40$ |
| Finetuned | $0.51$ | $0.49$ | $0.36$ | $0.51$ | $0.80$ | $0.78$ | $0.29$ |
| | | | OBTRUDE[b] | | | | |
| Frozen | $2.91$ | $3.78$ | $3.25$ | - | - | - | $1.41$ |
| Finetuned | $2.89$ | $3.68$ | $3.22$ | - | - | - | $1.38$ |

[a]Prediction RMSEs for one stratified holdout set and the SNase OBTRUDE test set. Frozen refers to the KaML-ESM2 model using frozen ESM2_650M embeddings. Finetuned refers to the KaML-ESM2 model finetuned using the parameter efficient QLoRA method.[15] No GAINES augmentation was applied in training.

Table S7: Performance of the structure-based KaML-CBT model with conventional and GAINES training[a]

| | | KaML-CBT[c] | | |
|---|---|---|---|---|
| | | RMSE | PCC | CER[b] |
| Asp | Conv | $0.75 \pm 0.04$ | $0.86 \pm 0.02$ | $13/916$ |
| | GAINES | $0.70 \pm 0.03$ | $0.89 \pm 0.01$ | $27/2052$ |
| Glu | Conv | $0.60 \pm 0.02$ | $0.84 \pm 0.01$ | $5/1076$ |
| | GAINES | $0.59 \pm 0.01$ | $0.86 \pm 0.01$ | $11/2388$ |
| Cys | Conv | $1.50 \pm 0.13$ | $0.61 \pm 0.12$ | $11/209$ |
| | GAINES | $1.01 \pm 0.09$ | $0.73 \pm 0.08$ | $4/147$ |
| Tyr | Conv | $1.24 \pm 0.19$ | $0.39 \pm 0.19$ | $3/31$ |
| | GAINES | $0.51 \pm 0.05$ | $0.08 \pm 0.12$ | $0/88$ |
| His | Conv | $0.85 \pm 0.03$ | $0.51 \pm 0.04$ | $11/209$ |
| | GAINES | $0.63 \pm 0.01$ | $0.75 \pm 0.02$ | $23/520$ |
| Lys | Conv | $0.70 \pm 0.05$ | $0.80 \pm 0.04$ | $1/325$ |
| | GAINES | $0.61 \pm 0.03$ | $0.86 \pm 0.02$ | $9/753$ |
| All | Conv | $0.77 \pm 0.02$ | $0.95 \pm 0.01$ | $44/2766$ |
| | GAINES | $0.66 \pm 0.01$ | $0.96 \pm 0.01$ | $74/65948$ |

Performance of the KaML-CBT model trained using either the conventional (Conv) or the GAINES protocol. The average and standard error of RMSEs, Pearson correlation coefficients (PCCs), and critical error rate (CER) across 50 holdout test sets are listed. CER refers to the fraction of misclassified protonation states at pH 7.[1] Metrics for the conventionally trained KaML-CBT are taken from Ref[1] across 20 holdout test sets.

Table S8: Ablations related to pretrained representations and task-specific learning

| | All | Asp | Glu | Cys | Tyr | His | Lys |
|---|---|---|---|---|---|---|---|
| 0-shot | $1.59 \pm 0.02$ | $1.80 \pm 0.03$ | $1.46 \pm 0.03$ | $2.96 \pm 0.15$ | $1.15 \pm 0.11$ | $1.25 \pm 0.03$ | $1.40 \pm 0.05$ |
| With task-specific learning | | | | | | | |
| 1-epoch | $0.90 \pm 0.02$ | $0.94 \pm 0.03$ | $0.70 \pm 0.02$ | $1.25 \pm 0.10$ | $0.72 \pm 0.05$ | $0.89 \pm 0.02$ | $1.18 \pm 0.04$ |
| Layer 0 | $1.22 \pm 0.02$ | $1.42 \pm 0.04$ | $1.07 \pm 0.02$ | $1.90 \pm 0.08$ | $0.60 \pm 0.05$ | $1.04 \pm 0.04$ | $1.11 \pm 0.03$ |
| Permutation | $1.37 \pm 0.03$ | $1.45 \pm 0.05$ | $1.20 \pm 0.04$ | $2.71 \pm 0.13$ | $0.92 \pm 0.09$ | $1.34 \pm 0.06$ | $1.33 \pm 0.06$ |
| Other protein LLM | | | | | | | |
| ProstT5[18] | $0.89 \pm 0.01$ | $0.93 \pm 0.2$ | $0.55 \pm 0.01$ | $1.86 \pm 0.08$ | $0.98 \pm 0.04$ | $1.12 \pm 0.01$ | $0.77 \pm 0.02$ |
| Baseline | $0.75 \pm 0.02$ | $0.68 \pm 0.04$ | $0.62 \pm 0.03$ | $1.23 \pm 0.11$ | $1.58 \pm 0.17$ | $0.81 \pm 0.05$ | $0.71 \pm 0.07$ |

Overall and amino acid-specific RMSE±standard error of the p$K_a$ predictions across 50 stratified hold-out sets. 0-shot: ESM2 predictions generated without performing any MLP training. 1-epoch: KaML-ESM2 task head trained for a single epoch. Layer 0: KaML-ESM2 task head trained using the first layer of ESM2. Permutation: KaML-ESM2 task head trained with the p$K_a$ labels randomly permuted. ProstT5:[18] an MLP task head on top of the protein LLM ProstT5 trained with the same settings. Baseline: KaML-ESM2 **without pretraining or GAINES**. **Note: the baseline and zero-shot metrics here are different from Table 1, because models in Table 1 used pretraining.**

#### Supplementary Figures

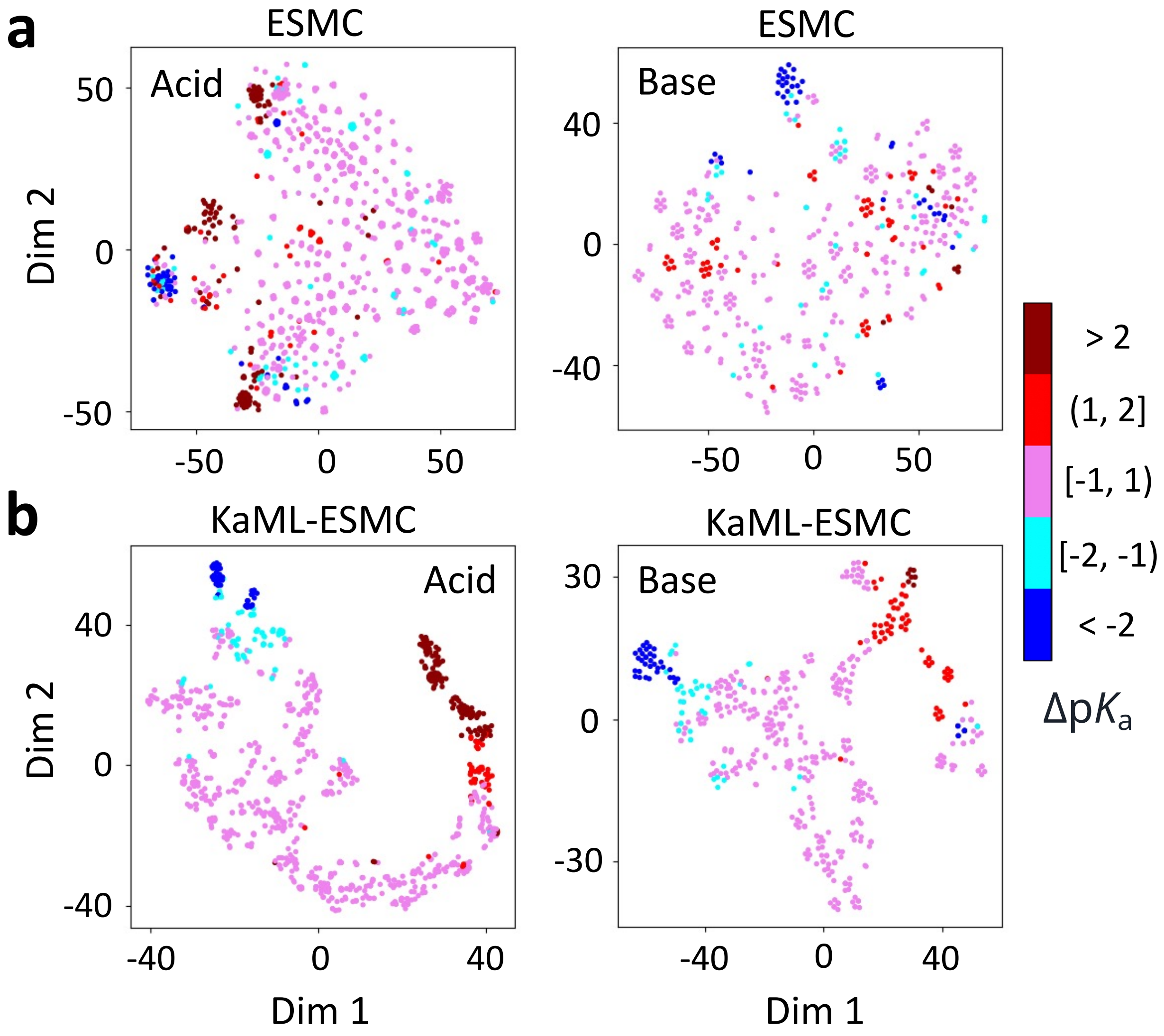

Figure S1: **Analysis of ESMC and KaML-ESMC representations in terms of amino acid identities and residue-specific p$K_a$ shifts.** t-SNE visualization of ESMC (top) and KaML-ESMC (last hidden layer, bottom) representations of acid (DECY) and base (HK) residues. Data points are colored according to the experimental p$K_a$ shifts relative to the model values. Embeddings were extracted from layer 80 of ESMC (2560-dimensional) and the last hidden layer of KaML-ESMC (64-dimensional). For the t-SNE algorithm, the maximum number of iterations is set to 1000, and the perplexity is set to 30 for each residue type.

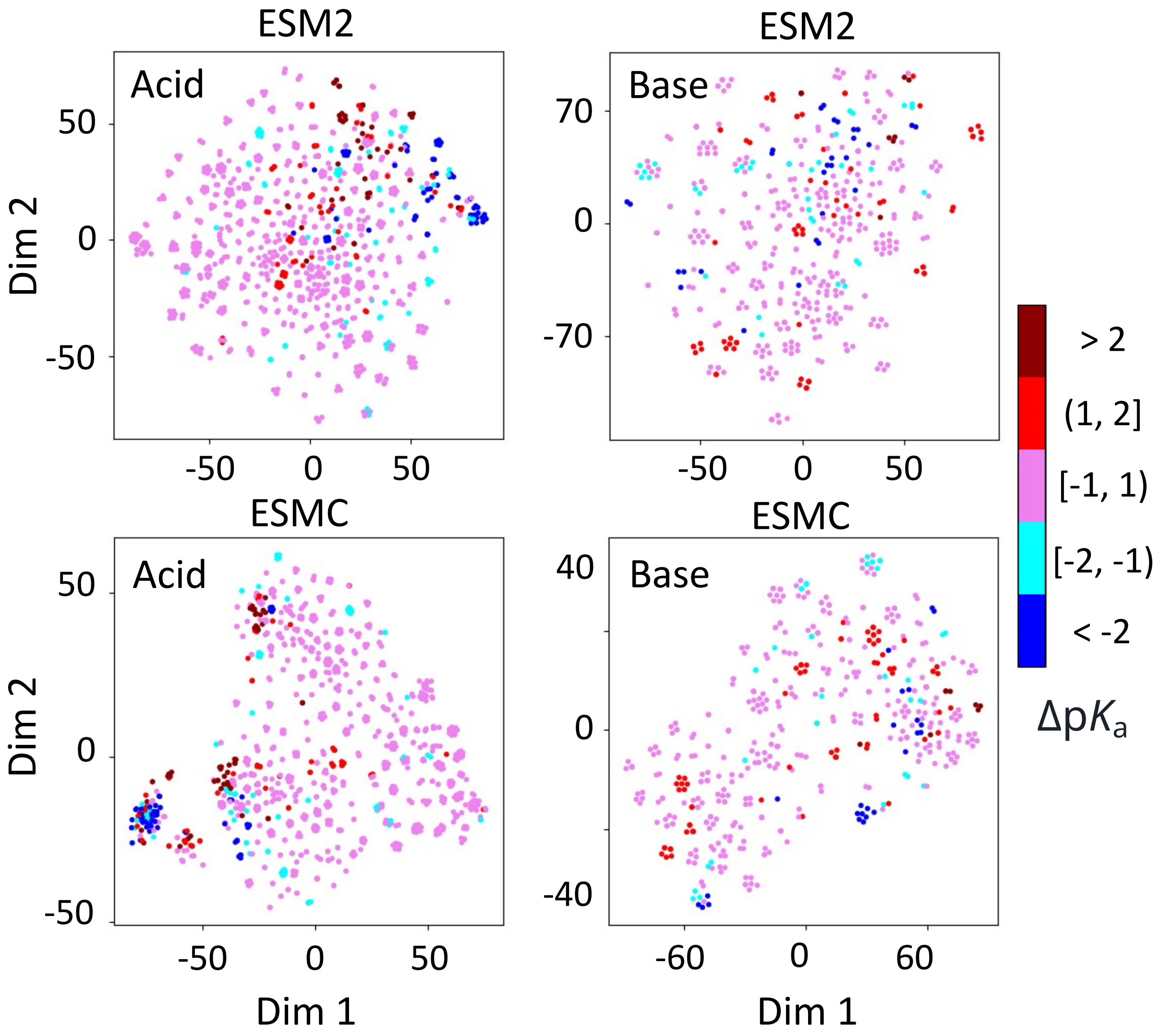

Figure S2: **t-SNE analysis of the ESM2 (top) and ESMC (bottom) representations of acidic and basic residues without the OBTRUDE set.** t-SNE visualization of the residue embeddings extracted from layer 31 of the ESMC model (1280-digit), and layer 80 of the ESMC model (2560-digit). For the t-SNE algorithm, the maximum number of iterations is set to 1000, and the perplexity is set to 30 for each residue type.

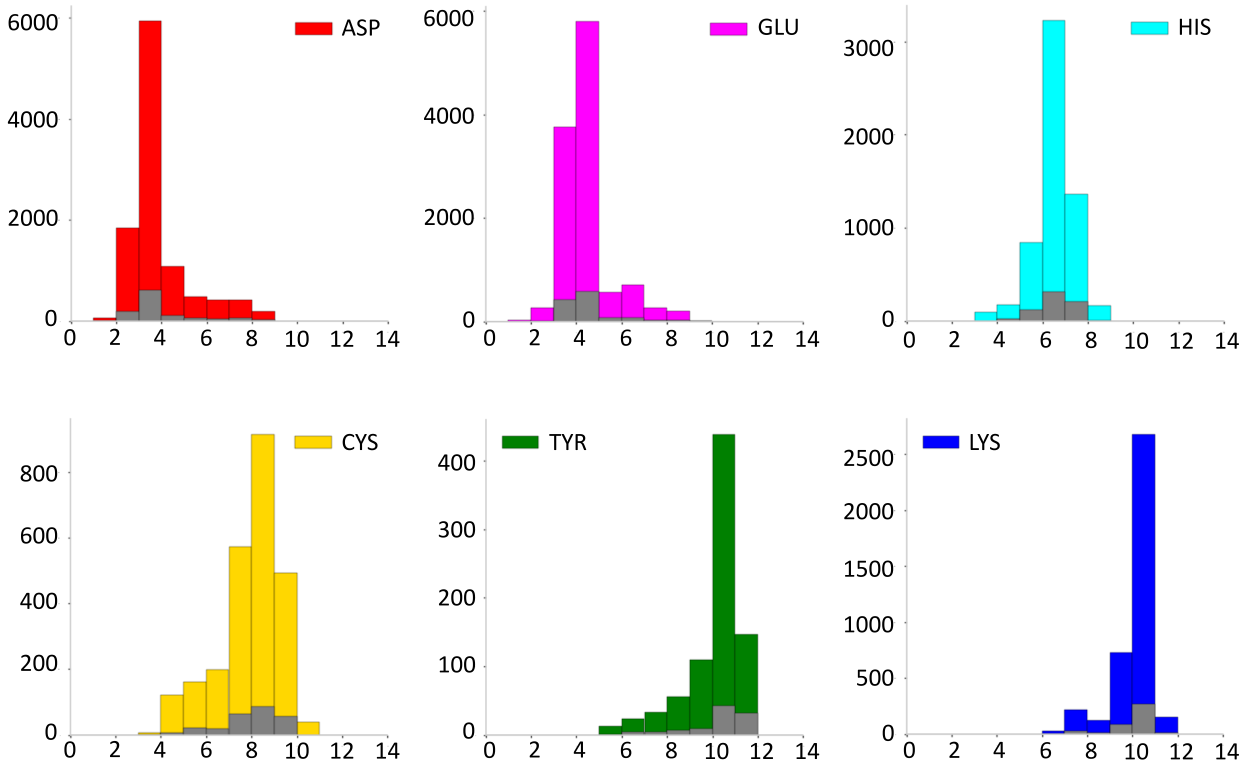

Figure S3: **Training-validation split for the pretraining dataset.** Histograms of the KaML-CBT predicted p$K_a$ values for pretraining KaML-ESM models. The dashed bins represent the validation dataset. KaML-CBT model taken from our previous work[1]

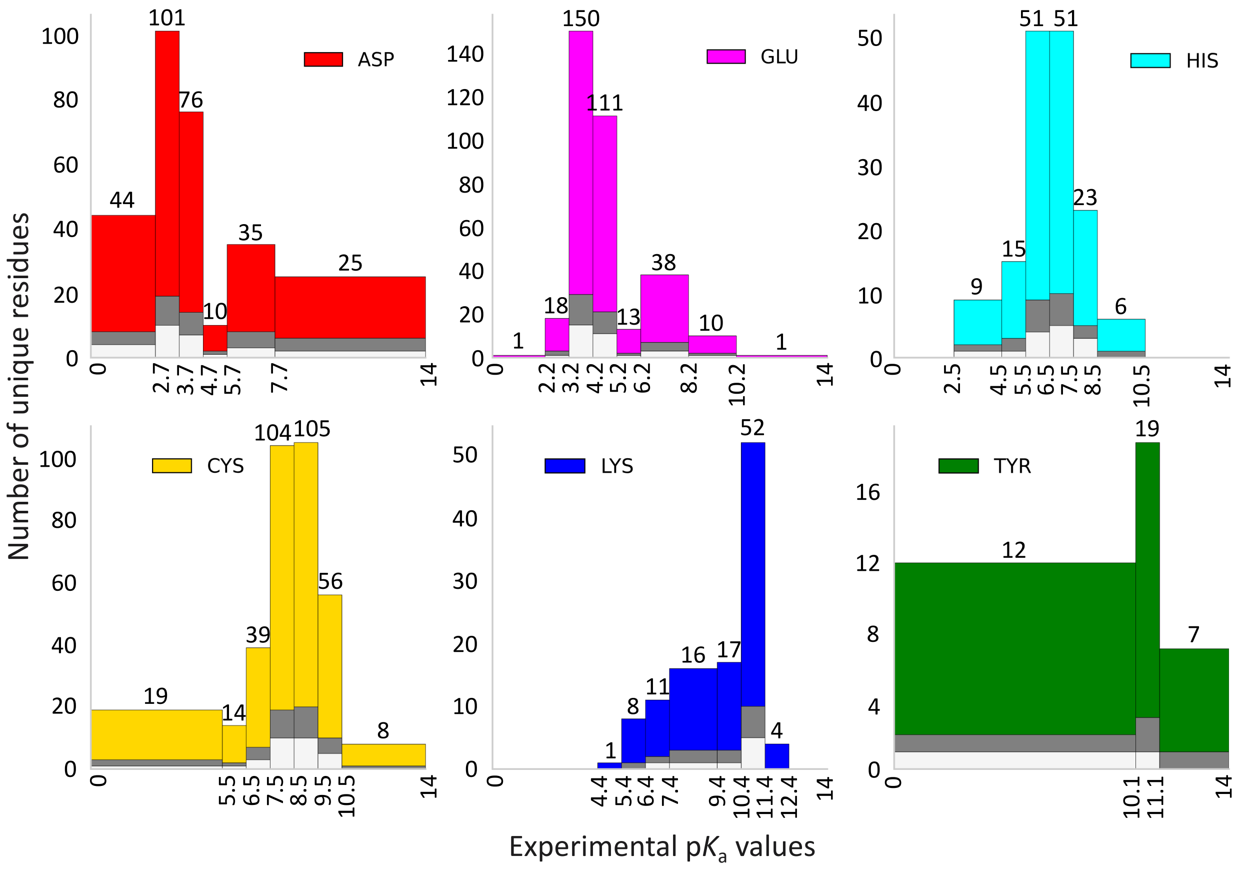

Figure S4: **Train-validation-test split for the PKAD-3 dataset.** The binning scheme used in this study for each residue type. The colored bins represent the overall dataset, gray bins represent the validation dataset, and the white bins represent the test dataset. The data splitting protocol is repeated 50 times independently, and the first splitting is shown here as an example.

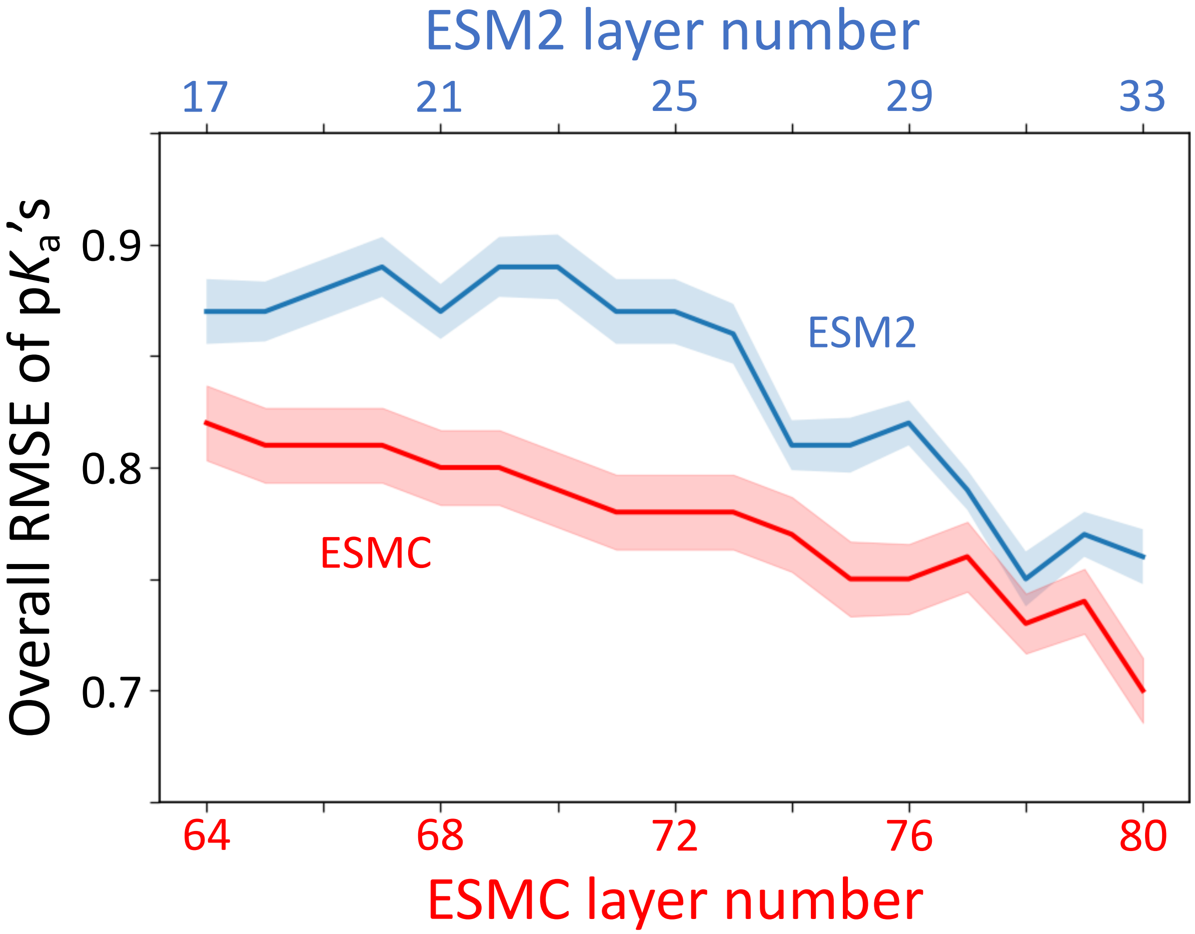

Figure S5: **ESM2 and ESMC exhibit distinct representation learning patterns across transformer layers.** Overall RMSEs of the p$K_a$'s predicted by models trained with embeddings from specific transformer layers up to the final layer (33 for ESM2_650M and 80 for ESMC_6B). The shaded regions represent the standard errors from 20 hold-out tests. Data for ESM2_650M and ESMC_6B are colored blue and red, respectively. No model pretraining was performed.

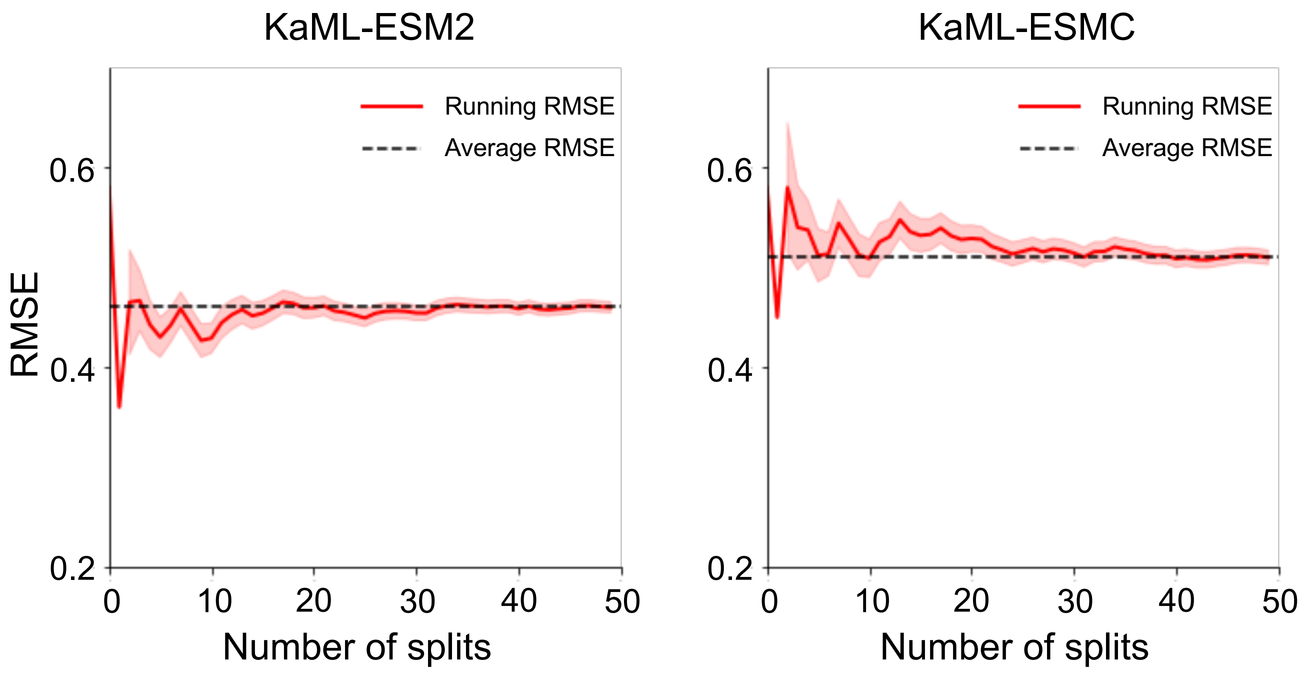

Figure S6: **RMSE convergence with respect to the number of random train-test data splits.** After 20 train-test splits, the running RMSE plateaus at 0.46 and 0.51, which is the average RMSE over 50 splits for KaML-ESM2 (left) and KaML-ESMC (right), respectively. Both models are trained with GAINES augmentation.

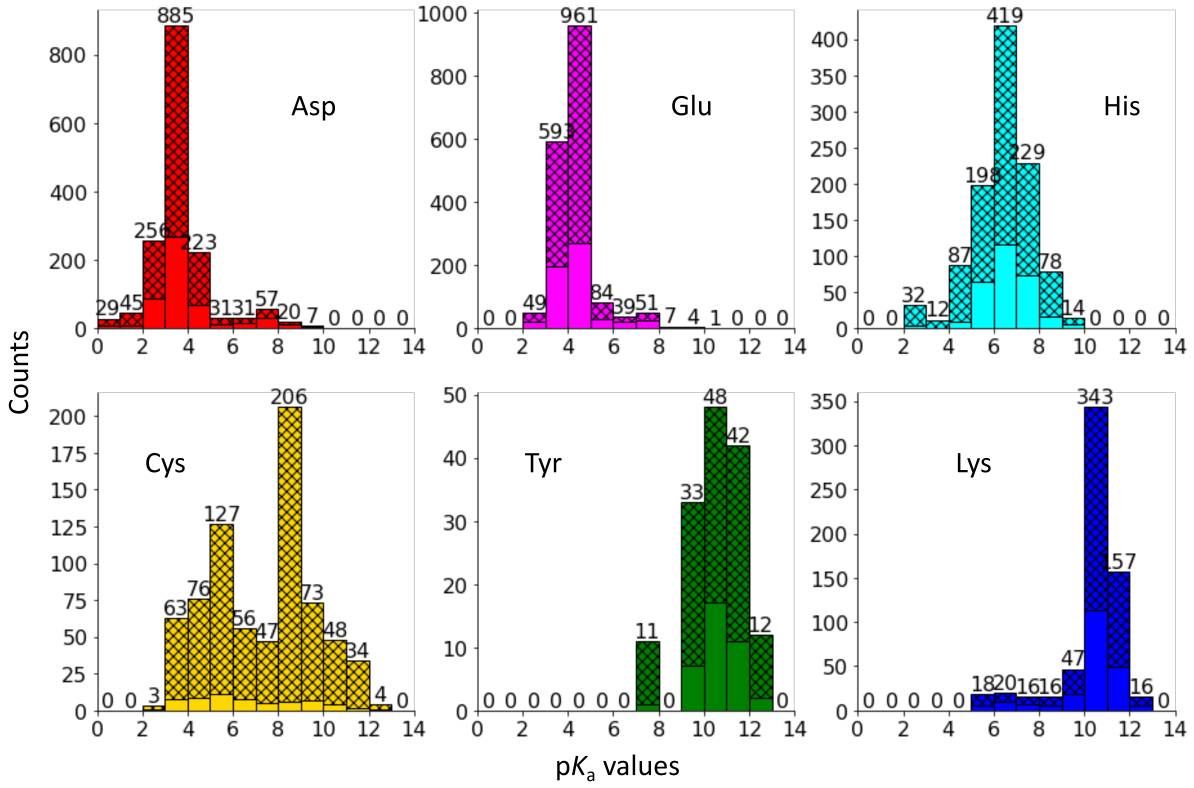

Figure S7: **Distribution of the GAINES augmented PKAD-3 dataset.** Histograms of the synthetic p$K_a$ values (shaded bins) and PKAD-3 experimental p$K_a$ values.

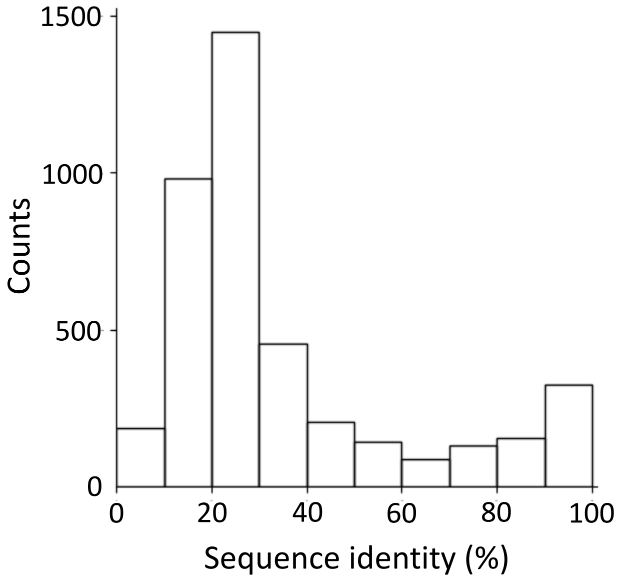

Figure S8: **Distribution of pairwise sequence identities between the query proteins in PKAD-3 and the corresponding value proteins.** Percent sequence identity is calculated between the query and value protein pairs.

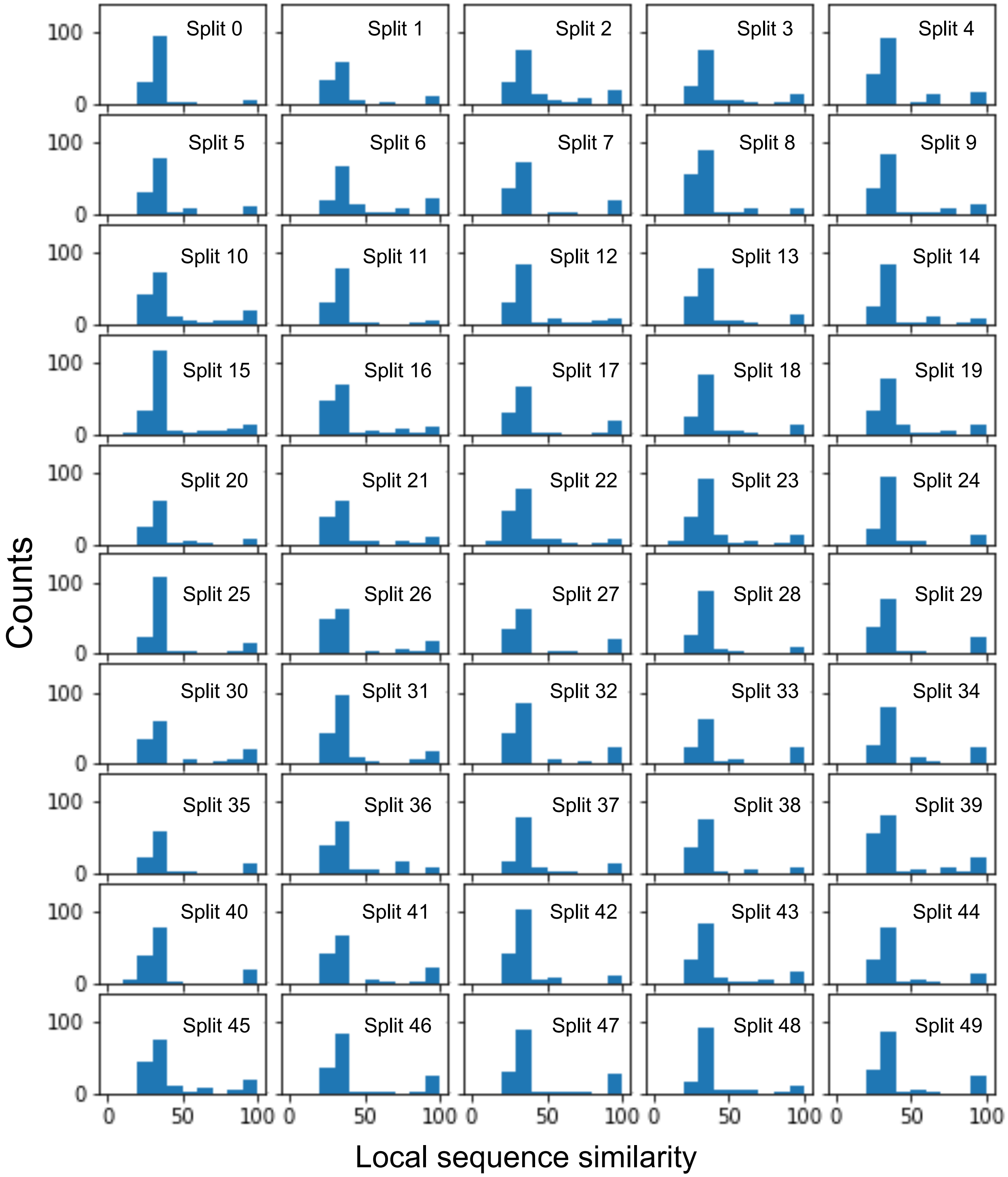

Figure S9: **Distributions of local sequence similarities between training and 50 hold-out sets (random training/holdout splitting).** For each residue in the test dataset, pairwise local sequence similarity was calculated against all residues in the training and validation datasets. The highest similarity score identified for each test residue was then recorded and plotted.

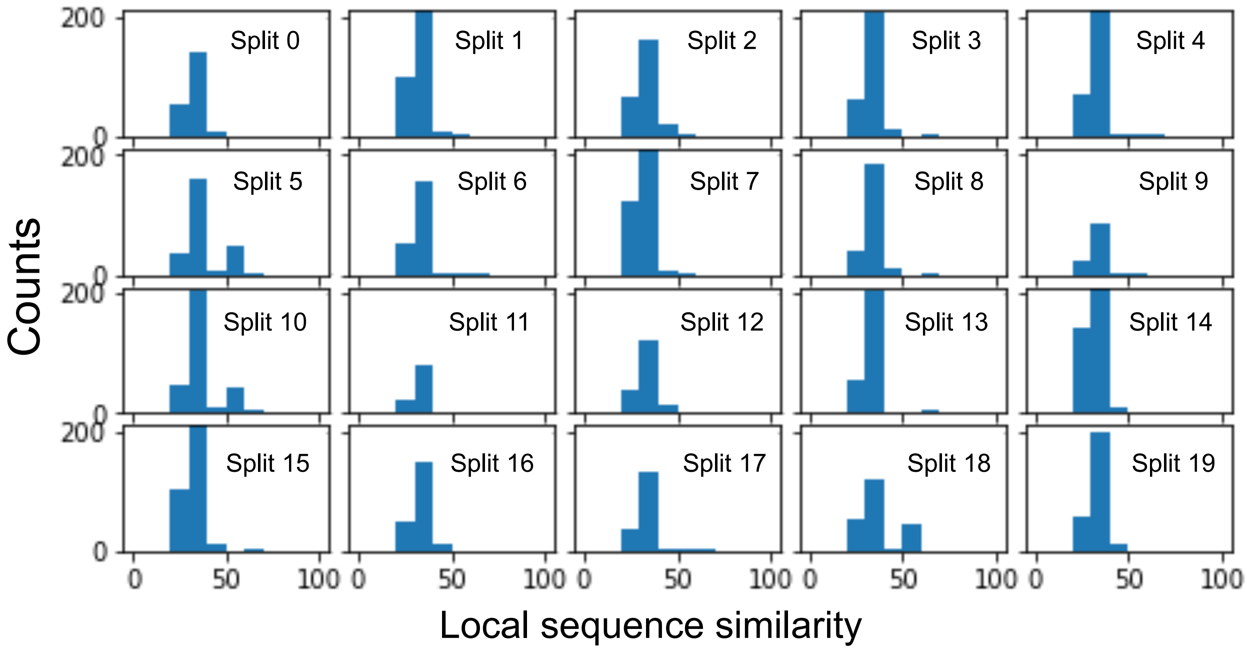

Figure S10: **Local sequence similarities between training and 20 random holdout sets (CD-HIT splitting).** For each residue in the test dataset, pairwise local sequence similarity was calculated against all residues in the training and validation datasets. The highest similarity score identified for each test residue was then recorded and plotted.

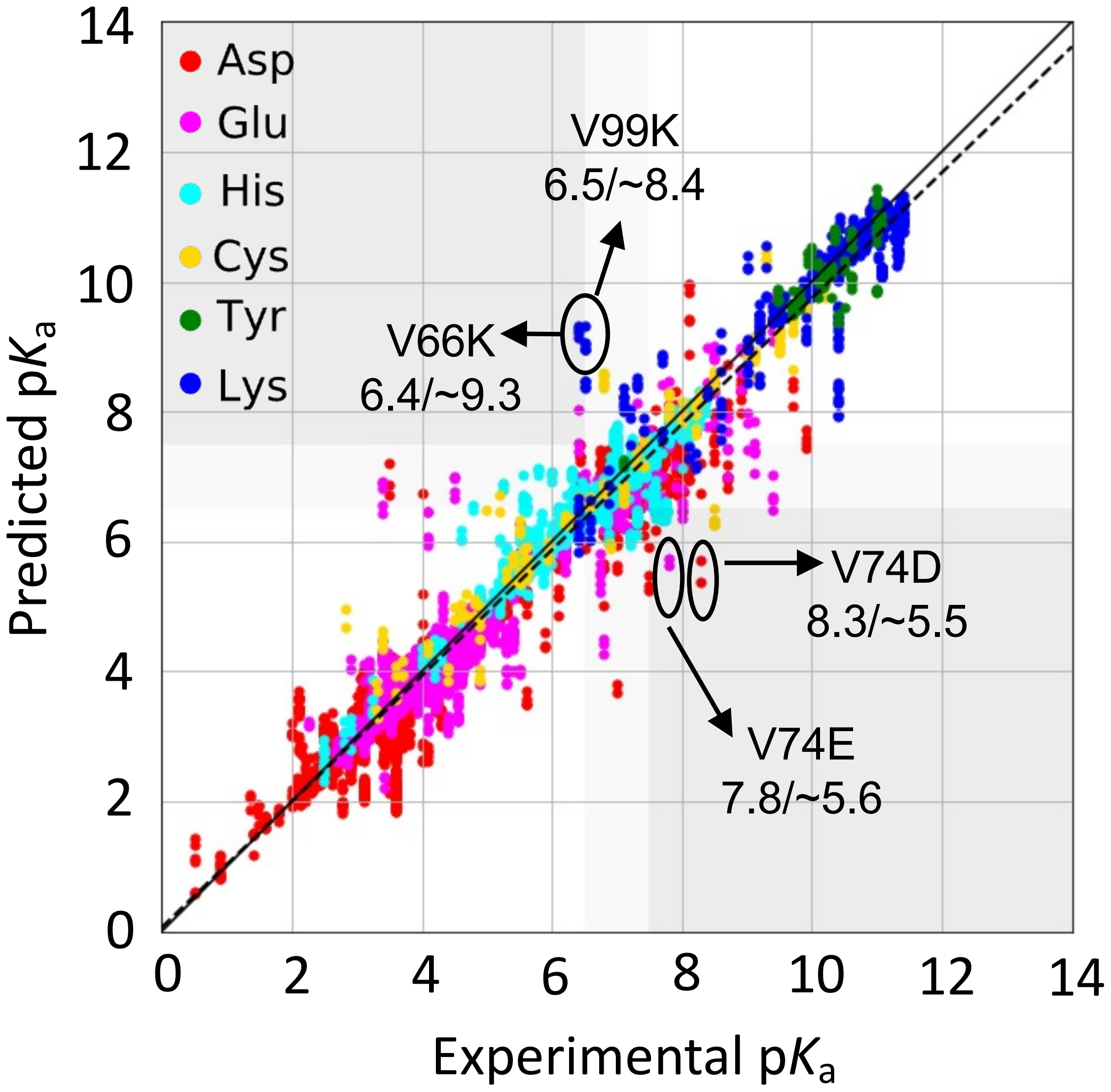

Figure S11: **Error analysis of KaML-ESM2 predictions across 50 holdout sets.** Experimental vs. KaML-ESM2 predicted p$K_a$'s across all 50 holdout sets. The solid line is the identity, and the dotted line is a linear fit. Data points are color-coded by amino acid. Following our previous work,[1] we divided the p$K_a$ range into different quadrants to illustrate whether a predicted p$K_a$ corresponds to the correct protonation state at pH 7 according to the experimental p$K_a$. White quadrants indicate correct protonation state predictions; gray regions indicate critical errors (i.e., protonated predicted as deprotonated or vice versa); and light gray regions indicate that the predicted p$K_a$ belongs to the titration range. Circled outliers are discussed in the main text.

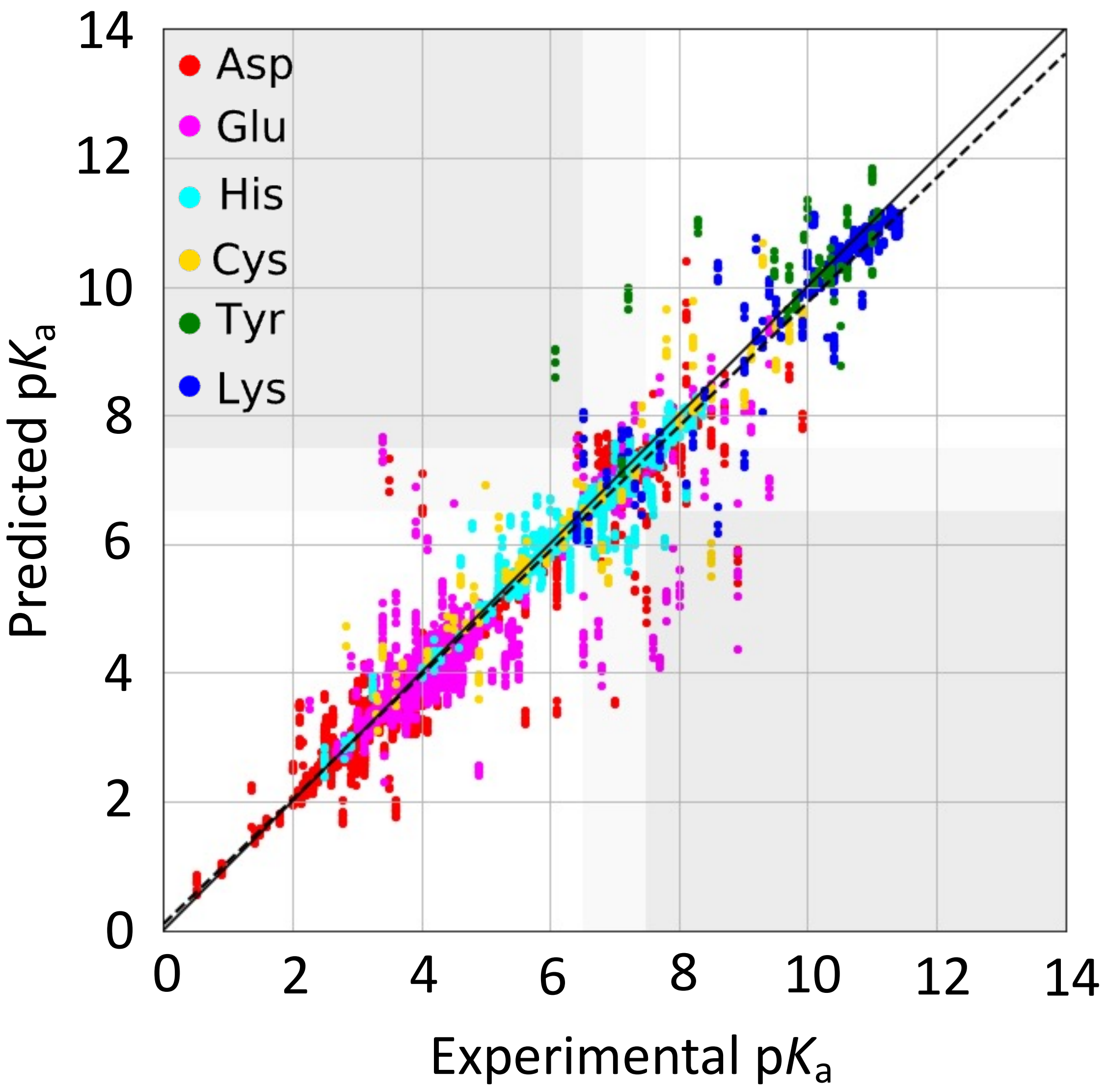

Figure S12: **Error analysis of KaML-ESMC predictions across 50 holdout sets.** Experimental vs. KaML-ESMC predicted p$K_a$'s in all 50 test sets. See Fig. S8 caption for detailed explanation.

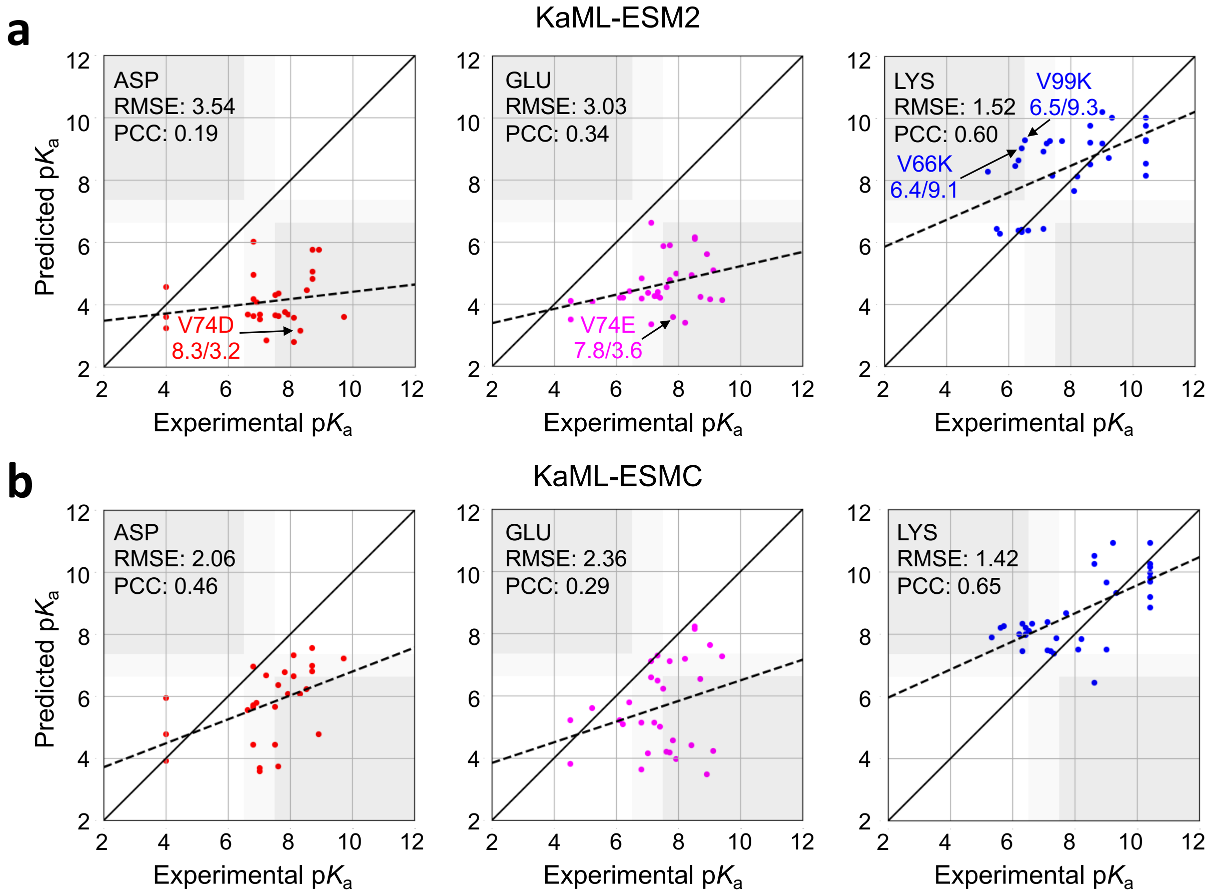

Figure S13: **Experimental vs. KaML-ESM2_no_OB (top) and KaML-ESMC_no_OB (bottom) predicted p$K_a$ values of the SNase OBTRUDEs.**

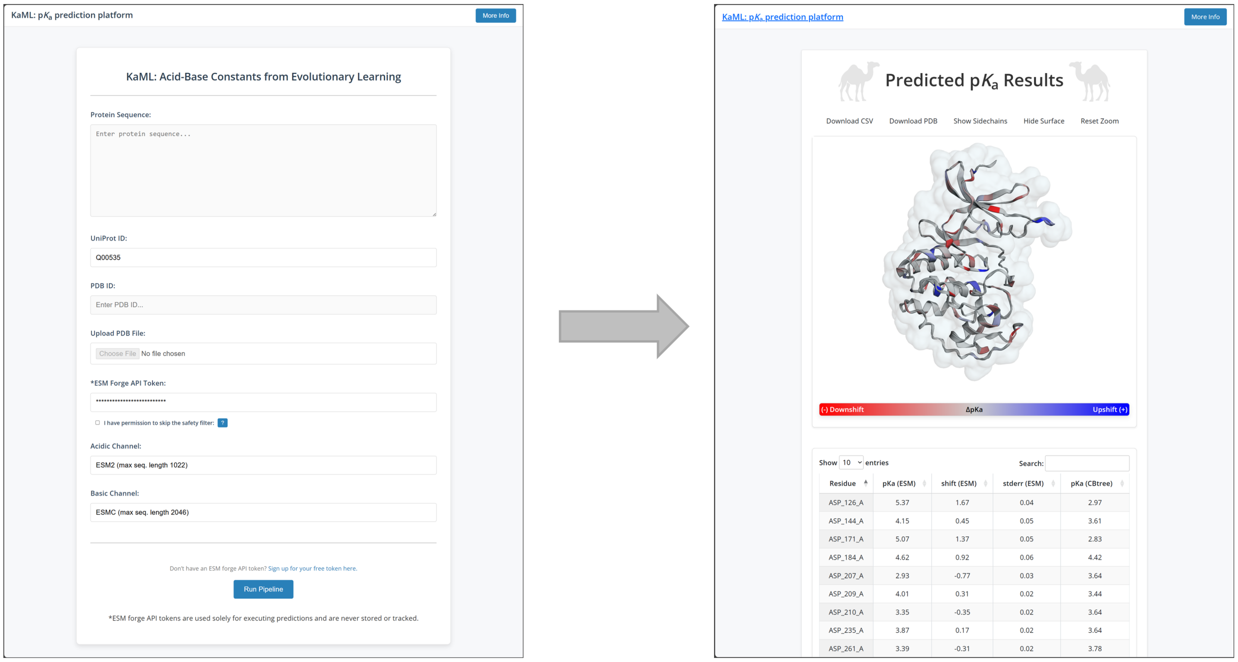

Figure S14: **Screenshots of the browser-based KaML platform.** An end-to-end web application (`kaml.computchem.org`) which takes either the protein sequence, Uniprot ID, PDB ID, or a PDB file as input to make p$K_a$ predictions. A command-line program is available at https: //github.com/JanaShenLab/KaML-ESM.

(27) Waterhouse, A.; Bertoni, M.; Bienert, S.; Studer, G.; Tauriello, G.; Gumienny, R.; Heer, F. T.; de Beer, T. A. P.; Rempfer, C.; Bordoli, L.; Lepore, R.; Schwede, T.

SWISS-MODEL: Homology Modelling of Protein Structures and Complexes. *Nucleic Acids Research* **2018**, *46*, W296–W303.
